# Static morphogen scaling enables proportional growth in tissue growth model inspired by axolotl limb regeneration

**DOI:** 10.1101/2025.02.14.638295

**Authors:** Natalia Lyubaykina, Dunja Knapp, Pietro Tardivo, Maximilian Kotz, Tatiana Sandoval-Guzmán, Benjamin M. Friedrich

## Abstract

Axolotls can regenerate lost limbs throughout life, while they continue to grow. This poses the question of how the size and pattern of a regenerating limb is matched to a widely varying animal size. Two interacting signaling molecules, SHH and FGF8, are produced at opposite sides of the regenerating limb and sustain tissue growth through a pair of oppositely-oriented signaling gradients. As the size of the regrowing tissue can vary more than three-fold depending on the size of the animal, it is unclear how the activities of these mutually dependent morphogens are maintained and subsequently terminated to determine appropriate growth. Scaling of limb regeneration suggests a size-dependent adaptation of morphogen gradient parameters. Inspired by this biological example, we theoretically investigate general mechanisms of morphogen-controlled growth arrest and proportional growth. In the proposed mechanism, tissue growth increases the spatial distance between the two morphogen gradients, which eventually arrests morphogen activity and growth. We put forward two distinct scaling scenarios of morphogen gradients: either dynamic scaling with blastema size, where morphogen gradient parameters change dynamically with the growing tissue, or static scaling with animal size, where morphogen gradient parameters stay constant during blastema growth and only depend on animal size. We show that static scaling ensures proportional growth, but dynamic scaling does not. We compare theory predictions to experimental quantification of SHH and FGF8 morphogen gradient parameters at different time-points of regeneration in different-sized animals, indicating static scaling for some morphogen parameters, which is sufficient to ensure proportional growth in our model.

---

Axolotls possess remarkable regenerative capabilities, including the ability to regenerate fully functional limbs even in adulthood [1]. Limb regeneration shares similarities with limb development [2, 3], which is largely conserved among vertebrates [4]. However, limb regeneration must also cope with the different sizes of juvenile and adult animals (Fig. 1A), which can differ up to ten-fold in linear size [5]. Regeneration to match animal size represents a general open problem of adaptive morphogenesis [6]. Limb regeneration in axolotls starts with wound closure after limb loss, followed by the formation of a bud called blastema consisting of undifferentiated progenitor cells, growth of this blastema, and eventually cell differentiation to give rise to various limb tissues (Fig. 1B). Blastema growth depends on animal size (Fig. 1C; wide-field images of corresponding stages, see Fig. S1). The result of the first phase of blastema growth is a hypometric limb (called “tiny limb” in [7]). This hypometric limb subsequently grows to its final size in a second growth phase, which is particularly pronounced in larger animals [7].

**FIG. 1.**
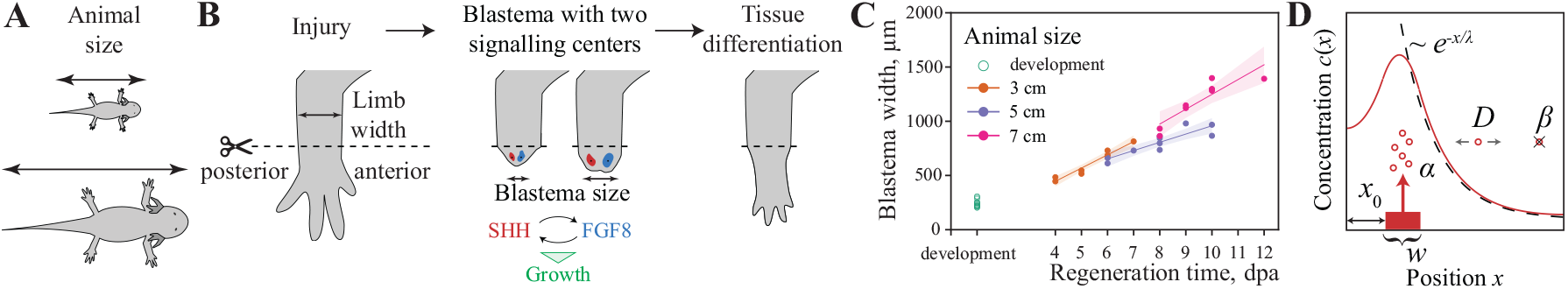
Axolotl limb regeneration. *A*. Schematic of axolotls of different sizes. *B*. Schematic of axolotl limb regeneration, highlighting its key stages: After limb amputation, a blastema with two distinct signaling centers producing SHH posteriorly (red) and FGF8 anteriorly (blue) forms. These mutually reinforcing morphogens induce cell proliferation, and thus regulate blastema growth. After this first growth phase of blastema growth, tissue differentiation and additional growth completes limb regeneration. Dashed line indicates amputation plane, arrows indicate limb and blastema size. *C*. Blastema size at subsequent time points (days post amputation, dpa) for animals of different size (filled symbols), and development (open symbols; measured along a line passing through the centers of SHH and FGF source regions in 3D images, see Methods for details). *D*. Morphogen gradient described by minimal reaction-diffusion model Eq. (1) together with visualization of model parameters.

We will focus on the first growth phase of limb regeneration, *blastema growth*. The sizes of same-stage blastemas scale with animal size [8]. This scaling is not perfectly proportional, but allometric, i.e., blastema size does not increase linearly with animal size, but according to a power-law with exponent 0.6 [8] (see also Fig. S2 in SI appendix). In contrast, intact limb size is proportional to animal size across a wide range of animal sizes (see also Fig. S3 in SI appendix). Thus, the relative size of the blastema is smaller in larger animals. For the smaller animals considered here, proportional scaling is an appropriate approximation.

Blastema growth involves a pair of oppositely-oriented morphogen gradients. Like the developing limb bud, the regenerating blastema display two signaling centers that secrete morphogens: a posterior center expressing SHH (Sonic Hedgehog) [9, 10], and an anterior center expressing FGF8 (Fibroblast Growth Factor) [11, 12], which mutually reinforce each other’s expression [13–15]. In intact mature limbs, SHH and FGF8 are undetectable [16, 17] (see also Fig. S4 in SI appendix). However, following injury, their production is activated, with clear evidence that both SHH and FGF8 are essential for maintaining blastema growth and successful limb regeneration [9, 11, 17]. Because of their mutual feedback, the individual roles of SHH and FGF8 in regulating cell proliferation are challenging to disentangle. While neither morphogen alone seems sufficient to support growth and regeneration in ectopically induced blastemas *in vivo* [17], both enhance proliferation individually and additively in cultured dissociated blastema cells *in vitro* [8]. Consequently, it remains challenging to determine whether one or both morphogens directly up-regulate proliferation.

Here, we focus on the stage of blastema growth as a paradigmatic model system for morphogen-dependent growth control and growth arrest in animals of different sizes. Recently, the sizes of the source regions of SHH and FGF8 were measured by chromogenic RNA *in situ* stainings in thin sections of middle-stage blastemas of animals of different sizes, indicating allometric scaling of source regions with animal size at the chosen time-point [8]. Inspired by this, we explore a minimal mathematical model of coupled tissue growth and morphogen dynamics in a system of two morphogens to identify physical mechanisms that ensure robust growth arrest and proportional growth. We systematically dissect differences between two fundamental growth rules based on either one or two morphogens, as well as two scaling scenarios of morphogen gradients, scaling either *dynamically* with growing blastema size, or *statically* with animal size. Our theoretical analysis singles out a mechanism of growth control by two oppositely-oriented morphogen gradients, where at least some morphogen gradient parameters must exhibit static scaling to enable proportional growth. By combining whole mount *in situ* hybridization with tissue clearing and confocal microscopy [18], we quantify the three-dimensional size of SHH and FGF8 morphogen sources as a function of time in regenerating blastemas of different-sized animals, as well as the FGF8 signaling range by measuring the expression of its down-stream target *Dusp6*. We found that some gradient parameters (FGF8 source size) scale *statically* with animal size, while others scale *dynamically* with blastema size. Such scaling is sufficient to ensure proportional growth in our model.

## RESULTS

### Coupling morphogen dynamics and growth

As a minimal model of morphogen dynamics in a growing blastema, we consider a one-dimensional system of size *L*, see Fig. 1D. We first introduce notation for the simple case of a single morphogen. The morphogen is produced within a source region of width *w* (located at a distance *x*_0_ from the tissue boundary), subsequently diffuses through the tissue with effective diffusion coefficient *D*, while undergoing degradation with degradation rate *β*. The dynamics of morphogen concentration *c*(*x, t*) as a function of time *t* is then given by

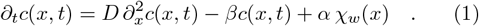

Here, the source function *χ*_*w*_(*x*) characterizes the shape of the morphogen source, which we assume to take the value 1 inside the interval [*x*_0_, *x*_0_ + *w*] and zero outside.

The rate total of morphogen production is thus *j* = *αw*, with units of molecules per time. The steady-state morphogen concentration profile *c*^*\**^(*x*) that is established at long times is thus fully defined by five parameters, which set its amplitude *A* and a characteristic pattern length-scale *λ*. This pattern length-scale is determined by a competition between the diffusion coefficient *D* and the degradation rate *β*

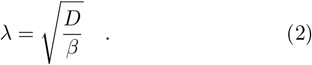

The steady-state concentration gradient *c*^*\**^(*x*) approximately follows an exponentially decaying profile with this pattern length-scale *λ*

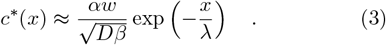

This approximation becomes exact when the morphogen source is small, localized on the system’s boundary, and the system is large. Analytic formulas for more realistic cases are given in SI appendix, Eq. (S18). Generally, concentrations are higher in smaller, bounded systems, as morphogens spread in a smaller domain. This size-dependence of morphogen concentrations will become important in our model, once morphogens control growth.

Eq. (1) generalizes in a straightforward manner to the case of two morphogens with respective concentration fields *s*(*x, t*) and *f* (*x, t*) instead of *c*(*x, t*) in Eq. (1), where *s*(*x*) and *f* (*x*) can be thought of as the concentrations of SHH and FGF8 morphogens. For conceptual clarity, we consider a symmetric case of two morphogens with identical kinetic parameters and mirror-symmetric source regions (see SI appendix, Eqs. (S24-25) for their formulas analogous to Eq. (1)). The general case of not-identical parameters for two morphogens is qualitatively similar.

To address the dynamics in growing tissue, we consider a one-dimensional tissue that grows with local growth rate *g*(*x, t*). Tissue growth convects the concentration field *c*(*x, t*) of morphogen with velocity *v*(*x, t*) where *∂*_*x*_*v* = *g*, as well as dilutes local morphogen concentration at rate *g* (see Methods).

### Growth control by one or two morphogens

Two interacting morphogens, SHH and FGF8, are implicated in regulating cell proliferation in axolotl limb development and regeneration [8, 17]. Due to the mutual feedback between these two morphogens, inactivating one consequently inactivates the other. Therefore, it is experimentally difficult to determine if the direct signal from just one or from both morphogens is needed to promote proliferation. To theoretically explore both cases, we define two putative growth rules, both of which are inspired by a switch-like control of cell behavior as a function of local morphogen concentration [19]. Specifically, we assume that the tissue grows locally if morphogen concentrations are above a threshold:

*One-morphogen growth rule:* the tissue grows at the position *x* if (and only if) the concentration *c*(*x*) of a morphogen is above a fixed threshold Θ, see Fig. 2A.

**FIG. 2.**
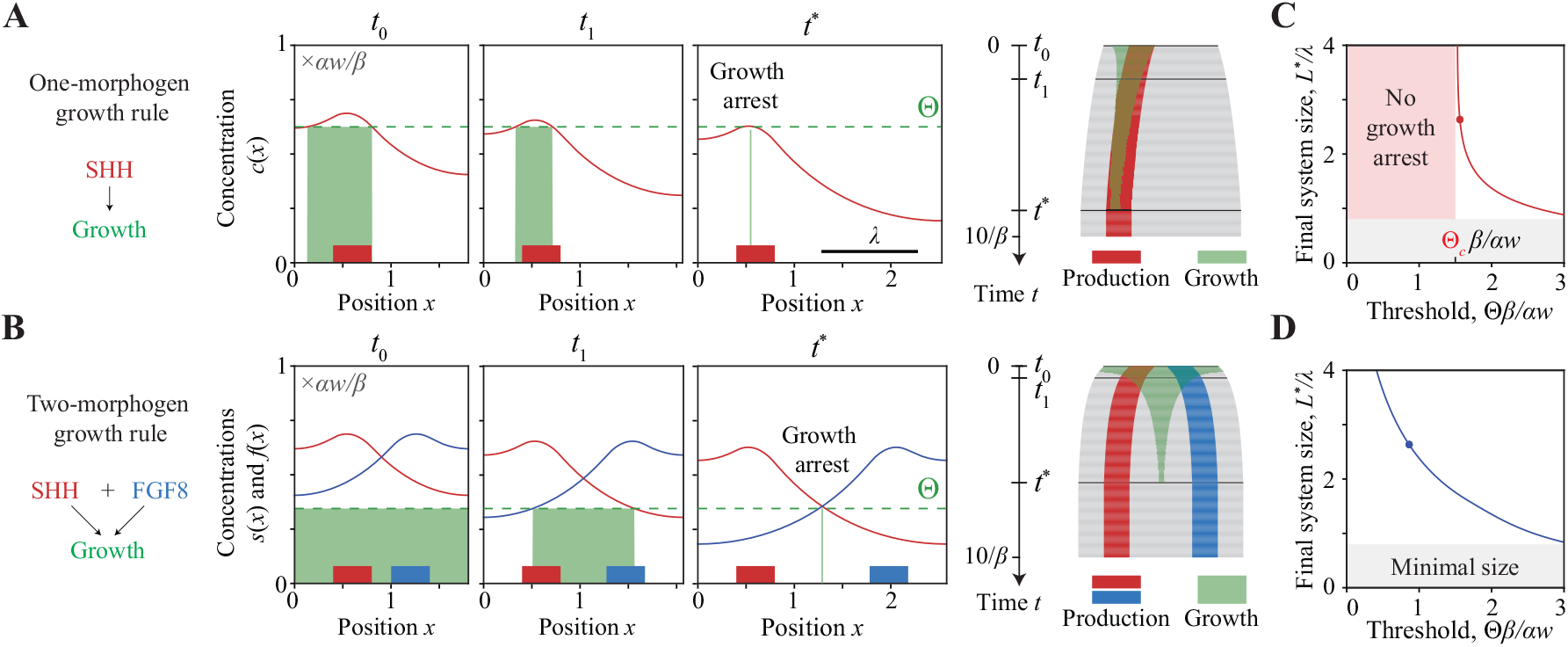
Two putative growth rules. *A*. Schematic of *one-morphogen growth rule*: the tissue grows where the concentration *c*(*x*) of a single morphogen is above the growth threshold Θ. Morphogen source region (red region), morphogen gradient (red curve), growth threshold Θ (dashed green line), and growth zone (green region) at different time points. A corresponding kymograph of simulated tissue growth until growth arrest is shown to the right (gray: tissue, red: morphogen source, green: growth zone with *c*(*x*) *>* Θ). *B*. Schematic of *two-morphogen growth rule*: a tissue grows only where the concentrations of both morphogens are above the growth threshold Θ. Morphogen gradients (red, blue) of the two morphogens at different time points, analogous to panel A. A kymograph shows production regions of both morphogens (red, blue). *C*. Final system size *L*^*\**^ normalized by pattern length-scale *λ* as a function of a normalized growth threshold Θ *β/αw* for the one-morphogen growth rule. Θ*c* indicates the critical threshold below which the one-morphogen growth rule fails to arrest growth. *D*. Final system size *L*^*\**^ normalized by pattern length-scale *λ* as a function of normalized growth threshold Θ *β/αw* for the two-morphogen growth rule. Threshold parameter and final system size in panels A and B are indicated by a point in panels C and D, respectively. Parameters: Table S1 in SI appendix; lengths are plotted relative to characteristic length *λ* = 1; concentrations are plotted relative to characteristic concentration *αw/β* = 0.4.

*Two-morphogen growth rule:* the tissue grows at the position *x* if the concentrations of both morphogens, *s*(*x*) and *f* (*x*) (e.g., SHH and FGF8) are above a fixed threshold Θ, see Fig. 2B.

For the sake of simplicity, we assume a symmetric scenario with equal threshold Θ for both *s*(*x*) and *f* (*x*) for the two-morphogen growth rule. A more general case can be mapped to this special case by a simple rescaling. Source width (*w*) and position (*x*_0_) are kept constant; more realistic scenarios of source growth are explored in the next section.

We numerically simulated morphogen dynamics according to Eq. (1) in a system with reflecting boundary conditions of dynamic size *L*(*t*), coupled with growth of this system assuming either the one- and two-morphogen growth rule, see Fig. 2. For both growth rules, the amplitudes of the morphogen gradients decrease as the system grows. Consequently, for the one-morphogen growth rule, the growth zone where morphogen levels exceed the threshold (*c*(*x, t*) *>* Θ) shrinks at its border opposite to the source, until – for suitable parameter choices – growth eventually arrests (Fig. 2A). For the two-morphogen growth rule, the growth zone where the levels of both morphogens exceed the threshold (*s*(*x, t*) *>* Θ and *f* (*x, t*) *>* Θ) shrinks even faster and at both borders, as the two opposing morphogen profiles overlap less when the system grows (Fig. 2B).

Fig. 2C and D show the final system size *L*^*\**^ at growth arrest for the one- and two-morphogen growth rule, respectively. For the one-morphogen growth rule, growth arrests only if the growth threshold Θ exceeds a critical value Θ_*c*_, where Θ_*c*_ = *αw/*(*Dβ*)^1*/*2^ for the limit case of a point source region (for details, see SI appendix, Eq. S28); for Θ *<* Θ_*c*_, the system will grow indefinitely (Fig. 2C). For the two-morphogen growth rule, growth always arrests for any threshold (Fig. 2D). Since we consider a source with constant size and position, and the source region should fit into the system, the size of the system should be at least *w* + *x*_0_, setting a minimal size. While we considered an idealized case where growth is slower than the dynamics of the morphogens, implying that morphogen gradients are always in quasi-steady state, numerical simulations reveal qualitatively similar behavior when this assumption is relaxed (SI appendix, Fig. S5). In the SI appendix, we also give analytic formulas for the final system size *L*^*\**^ as a function of the growth threshold Θ for the limit case of a point source region (Eqs. S27, S30). There, we also analyzed the sensitivity of the final system size *L*^*\**^ to the model parameters (Fig. S6), and found that the two-morphogen growth rule is less sensitive to parameter variations and ensures growth arrest for a wider range of parameters. Further, the two-morphogen growth rule arrests growth even for open or absorbing boundary conditions, while the one-morphogen growth rule requires reflecting boundary conditions (Fig. S10). This basic model shows that morphogen-dependent tissue growth can self-terminate, but does not include any scaling. Next, we investigate how tissues can attain the correct, proportionate final size through growth arrest in animals of different sizes.

### Dynamic vs. static scaling of morphogen gradients

In the generic model of growth control from the preceding section, the final system size *L*^*\**^ only depends on the initially defined morphogen gradient parameters. In axolotl limb regeneration, the final limb size depends on animal size, suggesting that some parameters of morphogen gradients should be size-dependent. In an idealized scenario of *proportional growth*, the final size *L*^*\**^ of the system should be proportional to animal size *L*_*a*_. Similarly, blastema size at the beginning of regeneration increases with animal size (Fig. 1C). Hence, in a simplified scenario, we assume that the initial system size *L*_0_ is proportional to animal size *L*_*a*_. To investigate how scaling of morphogen parameters could affect tissue growth, we investigate different scenarios for the size dependence of morphogen parameters.

We say that a morphogen gradient scales with a size *ℓ* if at least one of its features, source width *w* or pattern length-scale *λ*, both of which affect the signaling range, scales with this size *ℓ*. We distinguish two fundamentally different scaling scenarios, which differ by different interpretations of *ℓ*:

- *dynamic scaling* with growing blastema size *L*(*t*), corresponding to the choice *ℓ* = *L*(*t*), and
- *static scaling* with animal size *L*_*a*_, i.e. *ℓ* = *L*_*a*_.

For each of these two main scenarios, we consider three sub-scenarios. In all three sub-scenarios, we assume that the source region width and its position scale as *w ~ℓ* and *x*_0_ *~ ℓ*. In the first sub-scenario, which we term *source scaling*, the pattern length-scale *λ* is independent of size *ℓ*, see Fig. 3, left columns. In two additional sub-scenarios, in addition to source width, also the pattern length-scale scales as *λ ~ ℓ*. According to Eq. 2, either the effective diffusion coefficient *D* or the morphogen degradation rate *β* must change with *ℓ*. This defines a second and third sub-scenario, which we refer to as *D-scaling* (i.e., *D ~ ℓ*^2^), and *β-scaling* (*β ~ ℓ*^*−*2^), respectively, see Fig. 3, middle and right columns. The three sub-scenarios affect gradient amplitude *A* in different ways (SI appendix, Table S2): *A* increases with increasing size *ℓ* for source scaling, and even more so for *β*-scaling, where in addition to scaling of the source also the degradation rate *β* decreases for larger *ℓ*. In the case of *D*-scaling, the amplitude *A* stays approximately constant, as the effects of the source scaling and of the increase in diffusion cancel each other. Dynamic scaling has been reported in several systems [20–27], and shall therefore be analyzed first.

**FIG. 3.**
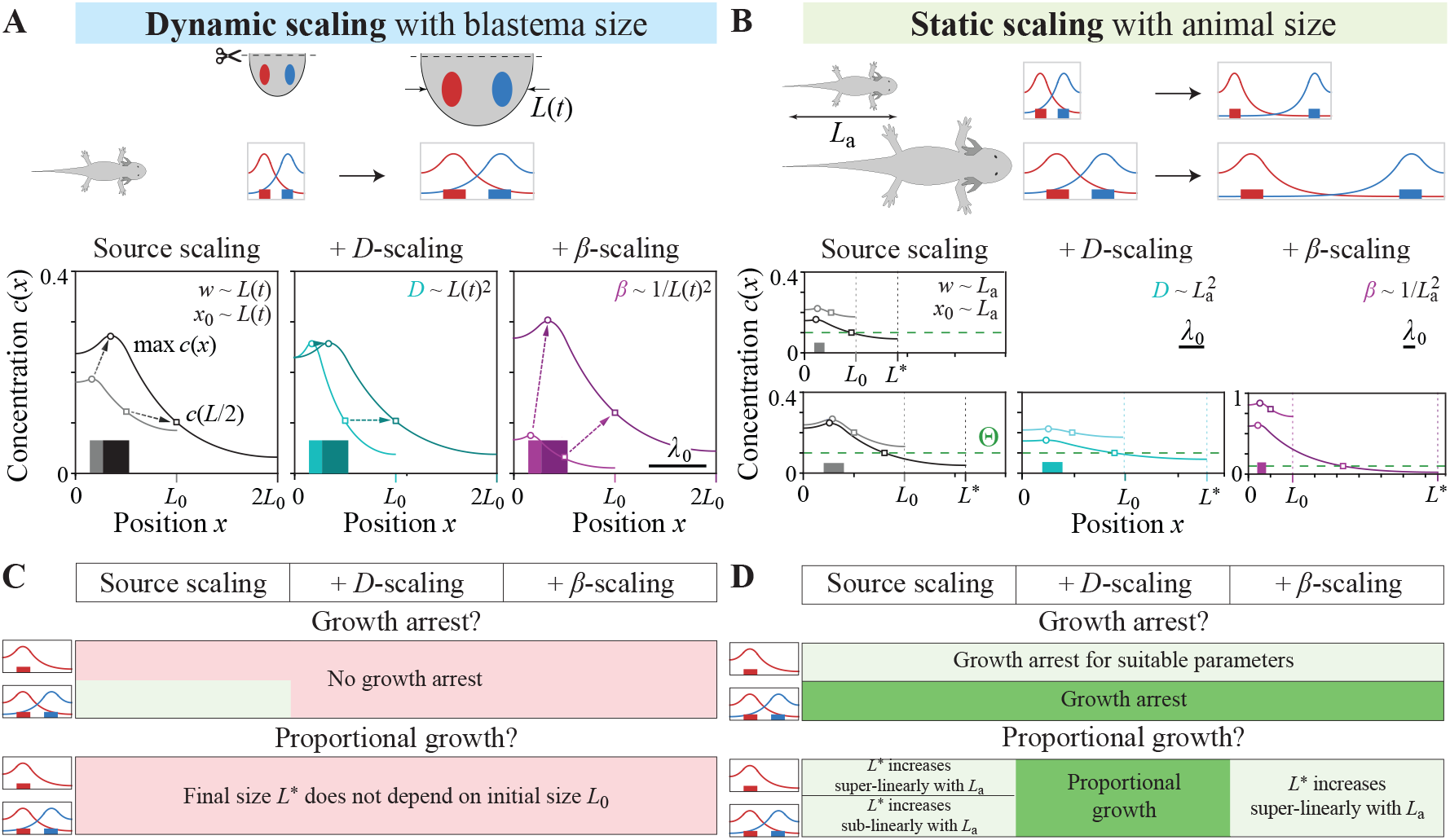
Two scenarios of morphogen gradient scaling. *A*. Schematic of *dynamic scaling* : morphogen gradients scale with growing blastema size *L*(*t*). Different sub-scenarios of dynamic scaling affect the amplitude of morphogen gradients differently (source scaling with production rate *α ~ L*(*t*): black, *D*-scaling with effective diffusion coefficient *D ~ L*(*t*)^2^: cyan, *β*-scaling with degradation rate *β ~* 1*/L*(*t*)^2^: magenta), see text for details. For each sub-scenario, source regions (boxes) and morphogen profiles (solid lines) are shown for initial tissue size *L*0 and when tissue size has doubled in lighter and darker color, respectively. For simplicity, the second morphogen is not shown; its profile is mirror-symmetric for the chosen symmetric parameters. For the one-morphogen growth rule, the system grows if the amplitude (circle) exceeds a threshold Θ; for the two-morphogen growth rule with symmetric morphogen profiles, the system grows if the concentration at the midpoint of the system (square) exceeds Θ. *B*. Schematic of *static scaling* with animal size: parameters of morphogen dynamics depend on animal size, but remain constant during growth. Same sub-scenarios of scaling as in panel A, but parameters of morphogen dynamics now depend on animal size *La*. Shown are morphogen gradients at the beginning of growth with initial system size *L*0 (assumed proportional to *La* for simplicity) and at growth arrest with final system size *L*^*\**^ for two values of *L*0 (upper and lower row). *C*. Summary of modeling predictions for *dynamic scaling* with blastema size, corresponding to the three scaling sub-scenarios introduced in panel A. For *dynamic scaling*, growth arrests for any growth threshold Θ for the two-morphogen growth rule if the morphogen source scales yet not the pattern length-scale *λ*, but not for any other sub-scenario. However, even in the scenario with growth arrest, the final system size *L*^*\**^ is independent of initial system size, i.e. there is no proportional growth. *D*. Summary of modeling predictions for *static scaling* with animal size, corresponding to the three scaling sub-scenarios introduced in panel B. For static scaling, morphogen dynamics parameters remain constant during growth. Correspondingly, growth arrests for any growth threshold Θ for the two-morphogen growth rule (introduced in Fig. 2), yet for the one-morphogen rule only if the growth threshold Θ exceeds a critical value Θ*c*. Proportional growth is achieved for the *D*-scaling sub-scenario, where both the effective diffusion coefficient *D* and the source region scales with animal size. Parameters: see Table S1.

#### a. Dynamic scaling with blastema size

In the absence of scaling, both the one-morphogen and the two-morphogen growth rules enable growth arrest. If the morphogen gradients exhibit dynamic scaling, model parameters and thus morphogen concentrations change dynamically with the growing blastema size *L*(*t*). For most scenarios of dynamic scaling, concentrations stay approximately constant or even increase, and hence growth does not arrest anymore, see Fig. 3A. The only exception is growth arrest for the special case when the pattern length-scale *λ* does not scale, and the two-morphogen rule is assumed. We obtain similar results without scaling of the source (SI appendix, Table S2). More generally, if either size *w* or position *x*_0_ of the morphogen source scale, we obtain a combinatorial set of 32 sub-scenarios: growth arrests in 10 of these sub-scenarios for the two-morphogen growth rule, but only 4 sub-scenarios for the one-morphogen growth rule (SI appendix, Figs. S7-S9). However, even if growth arrests, the final system size *L*^*\**^ is independent of the initial system size *L*_0_. Thus, in the case of dynamic scaling with growing blastema size, proportional growth is not possible.

#### b. Scaling with animal size

Next, we explore the scenario of scaling with animal size, where the parameters of the morphogen dynamics are set by the animal size *L*_*a*_ and then remain constant during blastema growth. This could occur if those parameters are set by systemic factors (e.g. hormonal or metabolic). As parameters of the morphogen dynamics remain constant during blastema growth, for animals of fixed size, the system behaves exactly the same way as in the case without scaling. We conclude that growth will arrest under minimal assumptions as described in Fig. 2. However, the final blastema size *L*^*\**^ now depends on the animal size *L*_*a*_ in a non-trivial manner, see Fig. 3D. Proportional growth with *L*^*\**^ *~ L*_*a*_
is obtained exactly for *D*-scaling. For *β*-scaling, *L* increases super-linearly with *L*_*a*_. These predictions are robust, and we obtain similar results, e.g., in the limit case of a point source (SI appendix, Table S2). While Fig. 3 focused on the limit case of slow growth *g ≪ β*, we observe virtually the same proportional growth for the two-morphogen growth rule (and variants thereof with graded growth response) if growth is fast (SI appendix, Fig. S5). Similarly, results virtually do not change if instead of the reflecting boundary conditions used in Fig. 3, we assume either open or absorbing boundary conditions, with the exception of the one-morphogen growth rule, which does not facilitate growth arrest for these alternative boundary conditions (SI appendix, Fig. S10). This, once more, highlights the robustness of the two-morphogen growth rule.

#### c. Combination of dynamic and static scaling

While Fig. 3C shows that dynamic scaling alone cannot ensure proportional growth, intriguingly, a combination of dynamic and static scaling can. For example, a scenario where all SHH parameters exhibit dynamic scaling and all FGF parameters exhibit static scaling ensures proportional growth over a five-fold range of initial system sizes if *D*-scaling (i.e., size-dependent regulation of effective diffusion coefficients) is assumed (SI appendix, Fig. S11). This remains true even if all morphogen parameters – except FGF source size – exhibit dynamic scaling. In fact, the structure of the mathematical model implies that if growth arrest occurs for any form of *D*-scaling, combining dynamic and static scaling of different morphogen parameters, growth must be proportional with *L*^*\**^ *~ L*_*a*_ (SI appendix).

### Quantification of size-dependent morphogen gradients

To assess putative scaling of morphogen gradients in regenerating axolotl limb blastema, we quantitatively analyzed SHH and FGF8 source regions in 3D in animals of different sizes and at several different regeneration time points. We used Hybridization Chain Reaction (HCR) RNA *in situ*, combined with tissue clearing and confocal microscopy to perform 3D quantitative measurements of SHH and FGF8 gradient parameters in developing limb buds and in regenerating limb blastemas from animals of three different sizes (snout-to-tail animal size: 3 cm, 5 cm, 7 cm). Measurements were taken at multiple time points (days post amputation, dpa) throughout the period when both signaling centers were present, see Fig. 4A. We first quantified blastema size along the line in 3D passing through SHH and FGF8 source region centers as in Fig. 1C, see Fig. 4B. Next, we quantified the linear sizes of SHH and FGF8 source regions by computing the cubic root of the volume enclosed by the 50%-iso-surfaces of the respective mRNA signals, which defines a lengthscale that characterizes source size. We also quantified the gap size *x*_0_ between the source regions and the tissue boundary corresponding to the source position. Finally, the spatial range of FGF8 signaling was inferred from the volume enclosed by the 50%-iso-surface of *Dusp6* mRNA, a well-established downstream target of FGF8 signaling [28–30]. All morphogen gradient parameters exhibit a clear size dependence. Yet, distinguishing dynamic and static scaling is complicated by the fact that blastema size and animal size are correlated (Fig. S2-S3). We first tested both a linear regression as a function of blastema size (left column in Fig 4D), and a linear regression as a function of animal size (right column). For both fit scenarios, we obtained satisfactory fits, with statistically significant size dependence, suggesting that both static and dynamic scaling may contribute to the observed size-dependent patterns. A contribution of dynamic scaling is suggested by the temporal increase of gradient parameters within some animal size groups, see SI appendix, Fig. S12. On the other hand, static scaling with animal size is compatible with the data, with *R*^2^ values only slightly smaller or even larger than those of the linear regression against blastema size. To gain further insight, we performed a partial correlation analysis on this data, which allows testing for conditional statistical independence (SI appendix, Table S3): After factoring out the influence of animal size, less than 0.1% of the remaining variance of FGF8 source width *w* could be explained by blastema size, with no statistically significant correlation. In contrast, an analogous analysis for SHH source size revealed a much higher explained variance of 58% with high significance levels *p <* 0.1%, indicative of a contribution of dynamic scaling for SHH. The statistical analysis for the other morphogen parameters suggests different contributions of static and dynamic scaling, see Fig. 4D and SI appendix. In conclusion, the data for the FGF8 source region and possibly other morphogen gradient parameters is compatible with static scaling. Note that in our model, it is sufficient if only a subset of morphogen parameters, e.g. only FGF8 source size, exhibit static but not dynamic scaling, to ensure growth arrest and proportional growth (SI appendix, Fig. S11).

**FIG. 4.**
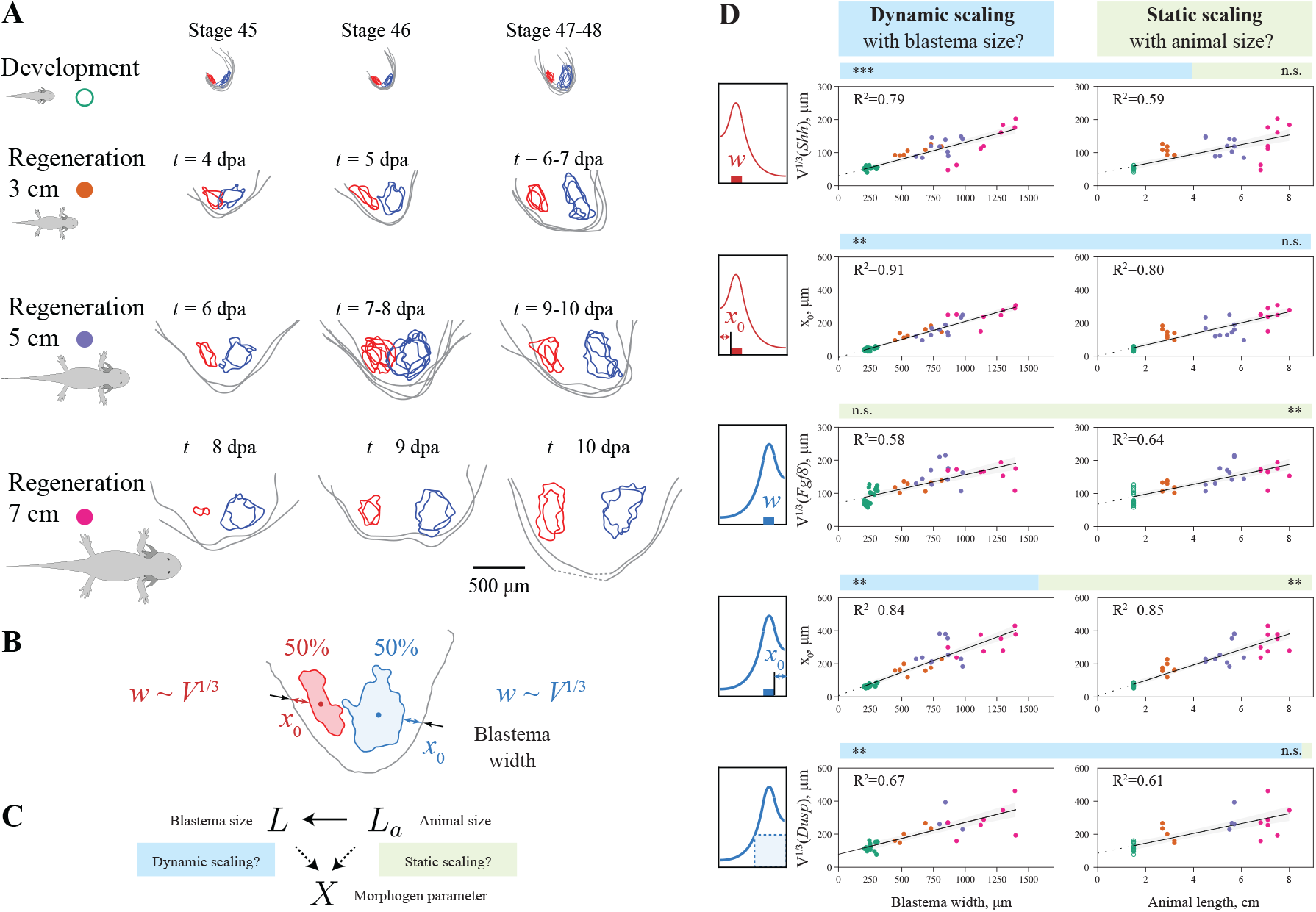
Size-dependence of SHH and FGF8 morphogen gradients. *A*. Outlines of SHH and FGF8 source regions from *in situ* hybridization in limb buds at different developmental stages, as well as regenerating blastemas of different-sized animals and at different regeneration time points (*Shh*: red, *Fgf8* : blue, 50% threshold of normalized intensity, shown as binary projections along dorsal-ventral axis on plane containing the centers of the *Shh* and *Fgf8* regions, see SI appendix for details; by convention, italicized names, e.g. *Shh* and *Fgf8*, refer to mRNA, indicating morphogen production, while capitalized names, e.g. SHH and FGF8, refer to the corresponding proteins. *B*. Definition of morphogen parameters: the linear source size *V* ^1*/*3^ determined from 3D-volume *V* enclosed by 50%-iso-surface for *Shh* defines the source width *w* for SHH, the gap size between SHH source boundary and tissue boundary defines the source position *x*0 (measured in 2D-projections along a line connecting the centers of the *Shh* and *Fgf8* regions as shown in panel A); analogous definitions for *Fgf8*. We determined a proxy for the FGF8 signaling range as *V* ^1*/*3^ from the 3D-volume *V* enclosed by the 50%-iso-surface of the FGF8 target *Dusp6. C*. A morphogen parameter *X* may exhibit either *dynamic scaling* with blastema size *L*, or *static scaling* with animal size *La*. Since *L* and *La* are correlated, causal inference is needed to test these scenarios. *D*. Morphogen parameters as defined in panel B as a function of blastema width (left column) and animal length (right column). Colors represent animal size as in panel A and Fig. 1C (filled circles: regeneration, open green circles: development; linear regression: black). For each morphogen parameter, a bar plot represents the result of a partial correlation analysis (statistical significance and proportion of residual variance from a test for the respective opposite null hypothesis, indicative of dynamic scaling (blue, null hypothesis: only static scaling), and static scaling (green, null hypothesis: only dynamic scaling), see SI appendix for details).

### Mutual feedback between morphogens

Previous works suggested a mutual feedback between SHH and FGF production [13–15]. Above, we assumed that both morphogens are produced independently of each other and their source regions and production rates remain constant during tissue growth without specifying biological mechanisms of maintaining morphogen production. Here, we theoretically describe a possible minimal feedback scenario. We assume that there are regions competent for SHH or FGF8 production (shown as transparent regions in Fig. 5A). This assumption is supported by distinct properties of anterior and posterior blastema regions [16, 31–33]. We further assume that SHH and FGF8 are produced in their respective competent region if the opposite morphogen concentration is above a feedback threshold Θ_fb_. With such mutual feedback, the system can be in one of two limiting regimes, depending on the relative magnitude of the growth threshold Θ and the feedback threshold Θ_fb_. In the first regime, growth arrests because the growth threshold is reached first (green region in Fig. 5B). The system thus behaves similar to the case without feedback described above and arrests growth, while morphogen production in still ON. In the second regime, the feedback threshold is reached first (orange region in Fig. 5B). As a consequence, the source regions of morphogen production shrink rapidly, and morphogen production is switched OFF. Without morphogen production, also tissue growth arrests. In this second regime, the feedback between the two morphogens functions as a tissue-level ON/OFF switch for growth. While Fig. 5B shows results for the two-morphogen growth rule, the system behaves qualitatively similarly for the one-morphogen growth rule. Signaling delays of morphogen feedback would result in a gradual decrease of morphogen profile amplitudes, but otherwise similar behavior.

**FIG. 5.**
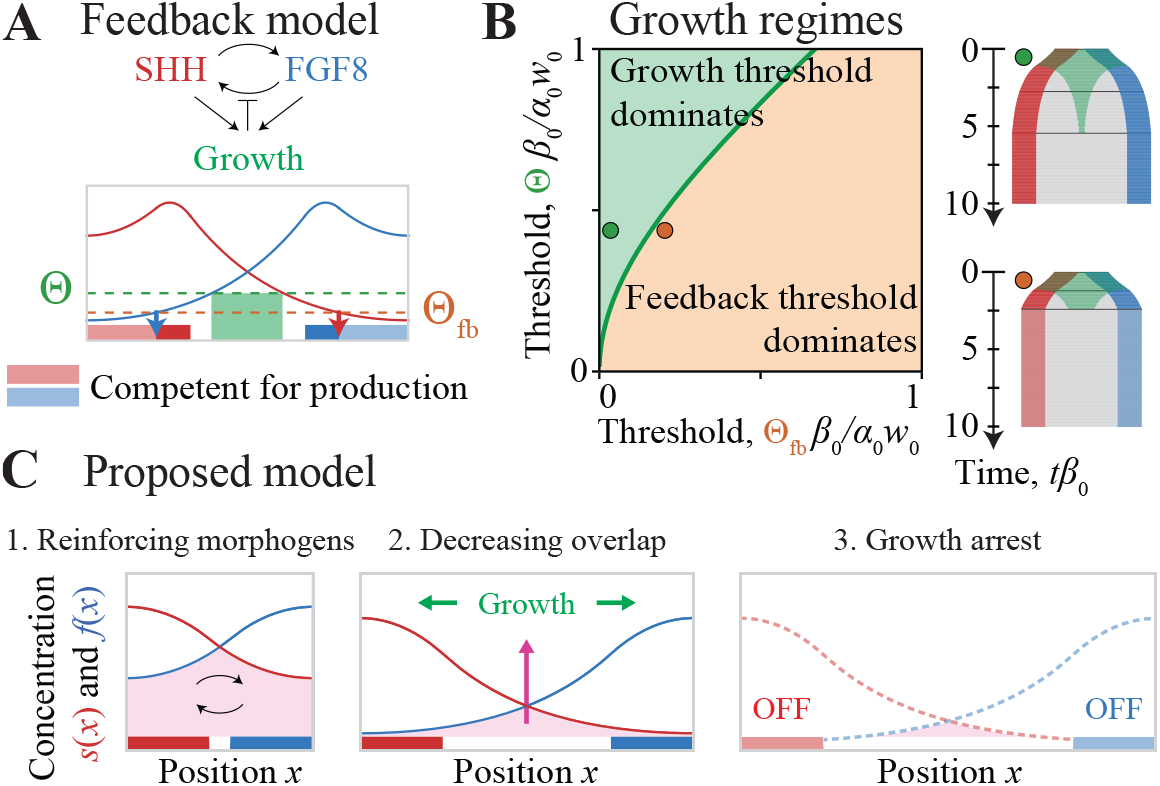
Feedback model of reinforcing morphogens. *A*. Minimal model of mutual feedback between oppositely-oriented morphogen gradients: Each morphogen can only be produced in a region of competent cells (transparent red and blue bars) where the concentration of the respective other morphogen exceeds a threshold (Θ_fb_, orange dashed), defining two source regions (solid red and blue bars). *B*. Phase diagram of growth regimes as a function of growth threshold Θ and feedback threshold Θ_fb_, here shown for the two-morphogen growth rule, exhibiting two regimes: the growth threshold Θ dominates, resulting in dynamics analogous to the case without feedback (green) as in Fig. 2, or, the feedback threshold Θ_fb_ dominates, resulting in instantaneous and homogeneous growth arrest across the entire system (orange). *C*. Model of growth arrest by the growth-threshold regime: 1. The tissue grows (green) where both morphogens (red and blue) are above the growth threshold (green dashed). 2. Tissue growth pushes the two morphogen sources apart, thus decreasing the region where the two morphogens overlap. 3. Due to the mutual feedback between the two morphogens, their respective source regions start shrinking rapidly, switching off both morphogens and arresting growth.

Adding feedback to the system thus does not significantly change its behavior with regard to growth arrest, demonstrating that our conclusions remain applicable also in a more realistic model. Feedback between the two morphogens could be a means of avoiding the slow fade-out of growth observed without feedback (Fig. 2), or even of switching off growth homogeneously throughout the system.

## DISCUSSION

Inspired by the size-adaptive regeneration of axolotl limbs, we addressed the general question of how morphogen-dependent tissue growth can be arrested once a tissue has reached its correct size. In our prototypical minimal model, tissue growth is induced by a threshold rule for either a single morphogen, or a pair of oppositely-oriented morphogen gradients. The two-morphogen growth rule with two oppositely-oriented gradients provides a particularly robust mechanism of growth arrest: the overlap between the two gradients induces growth; this growth increases the spatial separation between the two morphogen sources and thus decreases the overlap between the two gradients. This negative feedback loop robustly terminates growth at a specific tissue size. Proportional growth, however, where the final system size is proportional to initial size, requires scaling of morphogen gradients. We distinguished two fundamental scaling scenarios: *dynamic scaling* of morphogen gradients with the growing blastema, or *static scaling*, where morphogen gradient parameters depend on animal size, but remain constant during growth. Our theoretical analysis shows that static scaling with animal size of at least a subset of morphogen parameters is necessary and sufficient for proportional growth if a switch-like growth rule is assumed (Fig. 3 and Fig. S11 in SI appendix). We obtain similar results for graded growth responses (Fig. S5). Thus, different growth and feedback rules consistently yield robust growth arrest and proportional growth in the model. Yet, the spatio-temporal patterns of local tissue growth can differ (Fig. S13), which could help discriminating model variants in future experiments.

Blastema growth in axolotl regeneration is more complex. Our analysis addresses only the first stage of blastema outgrowth, which is followed by cell differentiation and a second growth phase. The second phase, which is independent of SHH-FGF signaling, is thought to be regulated by neuronal signals [7]. Nevertheless, the first growth phase and the blastema size at the onset of tissue differentiation is important to ensure accurate limb patterning, including the correct number of bone elements. Our model assumed reflecting boundary conditions along the anterior-posterior axis of SHH-FGF signaling. Indeed, the epidermis surrounding the blastema is expected to act as a diffusion barrier for spreading morphogens. However, morphogens may diffuse along the proximo-distal axis from the blastema into the limb stump at early stages, with ECM components possibly limiting morphogen outflux from the blastema, resulting in more complex boundary conditions. Reflecting boundary conditions near the source are important for the one-morphogen growth rule, while the two-morphogen growth rule works for different boundary conditions (Fig. S10).

Interestingly, blastema growth is not proportional to limb size in very large animals (≳ 10 cm), and is more accurately described by allometric scaling laws [8] (confirmed also in SI appendix, Fig. S2). This suggests limits of scaling. Note that sub-scenarios of our model can reproduce allometric scaling laws as observed experimentally (Fig. S5 in SI appendix). While our minimal model showed that exclusively dynamic scaling cannot enable proportional growth, it is likely that a combination of dynamic and static scaling applies in axolotl. Furthermore, dynamic scaling could still be compatible with proportional blastema growth if a separate timer mechanism terminates growth after a fixed time.

We used *in situ* hybridization in 3D tissues to quantify SHH and FGF8 source regions and FGF8 signaling range in developing limb buds and regenerating blastemas of different sizes at different time points. We find a clear size-dependence of morphogen gradient parameters. In particular, our data suggests static scaling with animal size for the FGF8 source region size, while the SHH source region size may exhibit dynamic scaling. This extends the recently published *in situ* hybridization analysis of source sizes in tissue slices for a single morphological stage in [8]. Our analysis of growth control by scaling morphogen gradients is independent of the particular implementation of morphogen scaling, and only depends on which parameters of morphogen dynamics are adjusted in a size-dependent manner. Scaling of morphogen gradients has been reported in a number of systems [20– 27]. A common concept in proposed mechanisms is the presence of a modulator such as an “expander”, whose concentration regulates, e.g., the effective diffusion coefficient or degradation rate of a morphogen [34–37]. In expander-dilution models, tissue growth dilutes the concentration of the expander inversely proportional to tissue volume [38]. In expander-repression models, the production of the expander depends on the morphogen itself [34]. Recently, SCUBE family molecules were shown to affect SHH spreading [39] and suggested to act by an expansion-repression mechanism to control the amplitude and pattern length-scale of the SHH gradient in the zebrafish neural tube [40], compatible with a mechanism of *D*-scaling.

SHH and FGF8 also guide limb development in other vertebrates [2, 3]. This pair of morphogen gradients must be switched on, but later also switched off again. Previous mathematical models focused on the temporal dynamics of morphogens [15], including the role of intermediate morphogens Bone Morphogenetic Protein (BMP) and Gremlin (GREM), which itself are thought to be a part of the SHH-FGF feedback loop. To explain how SHH and FGF become switched off again, various hypotheses reviewed in [41] were put forward: A first hypothesis suggests that the medial portion of the SHH competent region, which previously expressed SHH, loses this competence, creating an expanding region of cells that are not competent to express either SHH or GREM or FGF8 [42]. A second hypothesis suggests that FGF8 terminates GREM in a concentration-dependent manner [43]; alternatively, a low concentration of BMP could activate GREM at early stages, while a higher concentration of BMP could suppress GREM later [44]. We comment that these hypotheses would require signal delays to prevent becoming trapped in an intermediate state between “on” and “off”. Another hypothesis would be a timer mechanism that switches off SHH and FGF after a preset growth period [45]. These hypotheses not only underline the intricate molecular interactions involved in the growth control of developing and regenerating limbs but also start to appreciate the importance of spatial patterns of signaling.

Our model proposes that a coupling of spatial patterns and growth provides a simple and robust mechanism by which the SHH-FGF feedback loop could become switched off after the first stage of blastema outgrowth is completed (Fig. 5C). Our minimal model suggests specific experiments. Quantifying SHH and FGF8 morphogen gradients as a function of animal size and time, ideally by measuring protein concentrations instead of down-stream signaling, will allow testing our theory prediction of static scaling with animal size (or limb size). Cell culture experiments with controlled levels of morphogens as initiated in [8] could test the growth rules proposed here. This could pave the road towards more realistic, data-driven models that capitalize on the concepts laid out in the idealized, one-dimensional model studied here. In conclusion, we proposed a simple mechanism of two opposing morphogen gradients that reinforce each other, to enable robust growth arrest and proportional tissue growth by scaling of morphogen parameters with animal size.

## Methods

### Numerical methods

In the presence of tissue growth with local growth rate *g*(*x, t*), (1) changes to

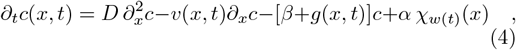

where *∂*_*x*_*v* = *g* [21, 46]. This equation is numerically solved using a modified Euler scheme in a Lagrangian frame with non-uniform spatial grid, whose bin sizes Δ*x*_*i*_ are updated according to Δ*x*_*i*_(*t*_*n*+1_) = [1 + *g*(*x*_*i*_, *t*_*n*_)Δ*t*]Δ*x*_*i*_(*t*_*n*_).

*Whole-mount HCR in situ hybridization and imaging Shh, Fgf8* and *Dusp6* mRNAs were detected in formaldehyde-fixed limb blastema and developing limb buds obtained from different-sized animals at different timepoints by whole mount HCR, a fluorescent RNA *in situ* method as described in [47] with minor modifications, followed by refractive index matching and confocal imaging in 3D (for detailed protocol, see SI appendix).

### Image analysis

For analysis of 3D multi-channel z-stack images, blastemas were segmented using APOC [48] trained with user-specified ground truth labels applied to the *Shh* channel using Napari [49] and custom Python code. Blood vessels detected in the autofluorescence channel were segmented using APOC, and the corresponding binary mask was excluded from all channels. The linear size of SHH and FGF8 source regions, as well as of FGF8 signaling indicated by *Dusp6* mRNA, were determined as the cubic roots of the volumes enclosed by the respective 50%-iso-intensity surfaces (after denoising and min-max normalization of each channel). Blastema size was measured in 3D images along a line passing through the centers of the *Shh* and *Fgf8* regions. Gap sizes between the blastema boundary and the boundaries of these source regions (*x*_0_) were measured along the same line, yet using 2D-projections to account for a possible non-convex shape of source regions.

### Data, Materials and Software Availability

Raw image data and Python code for growth simulations and image analysis can be accessed at a public github repository at https://github.com/NataliaLyubaykina, folder axolotl limb regeneration s. All other data, methods and analyses are included in the main text or in SI appendix.

## ACKNOWLEDGMENTS

We thank Leo Otsuki and Elly M. Tanaka (IMBA) for the transgenic axolotl line (tgSceI(ZRS:TFPnls-T2A-ERT2-Cre-ERT2)^Etnka^. We thank Elly M. Tanaka, as well as all members of the Sandoval-Guzmán and Friedrich groups for stimulating discussions, the animal carers of the CRTD (Beate Gruhl and Anja Wagner) for dedicated axolotl care, the light microscopy facility of the CMCB for excellent support, as well as Veikko F. Geyer for statistical analysis support. TSG and BMF acknowledge funding from a joint D-A-CH grant from the German and Austrian Science Foundations (441649267, jointly with Elly M. Tanaka). BMF was additionally supported by the Deutsche Forschungsgemeinschaft (DFG, German Research Foundation) under Germany’s Excellence Strategy - EXC-2068-390729961, as well as through a Heisenberg grant (421143374).

