## Supporting Information for "Static morphogen scaling enables proportional growth in tissue growth model inspired by axolotl limb regeneration"

This PDF file includes:

1. Figures S1 to S13
2. Tables S1 to S3
3. Supplementary Methods
4. Supplementary Information for Mathematical Modeling

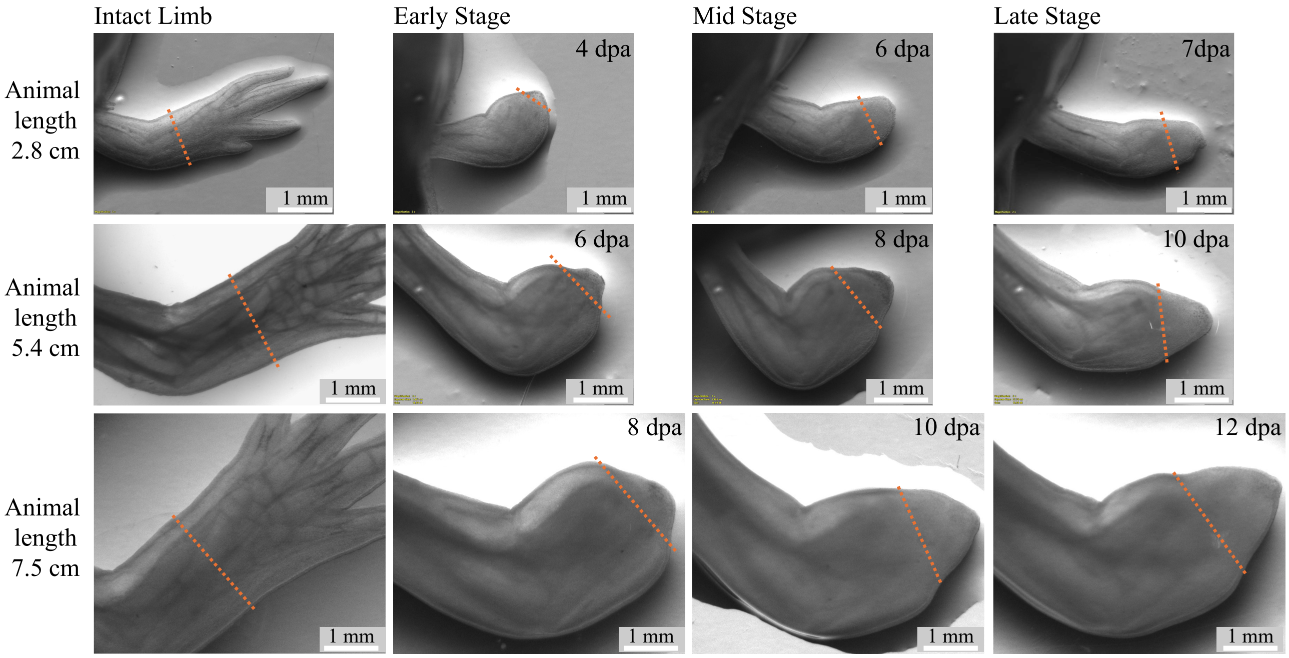


FIG. S1. *Regeneration time-course in animals of three different sizes as used for the quantification of morphogen parameters.* Intact limbs and three analogous stages of regeneration (early, middle and late blastema stages) are shown for animals of different sizes: 2.8 cm, 5.4 cm and 7.5 cm length from snout to tail. Analogous time points were designated based on morphology and SHH and FGF8 morphogen dynamics. The orange dotted lines indicate the amputation plane. Days post amputation (dpa) are indicated. Early stage: initial cohort of undifferentiated cells accumulated underneath the wound epidermis and onset of SHH and FGF8 expression, mid-stage: fast-growing blastema, FGF8 expressed in a single contiguous cloud, late stage: blastema began to flatten along the dorso-ventral axis, FGF8 source region begins to split into dorsal and ventral cloud.

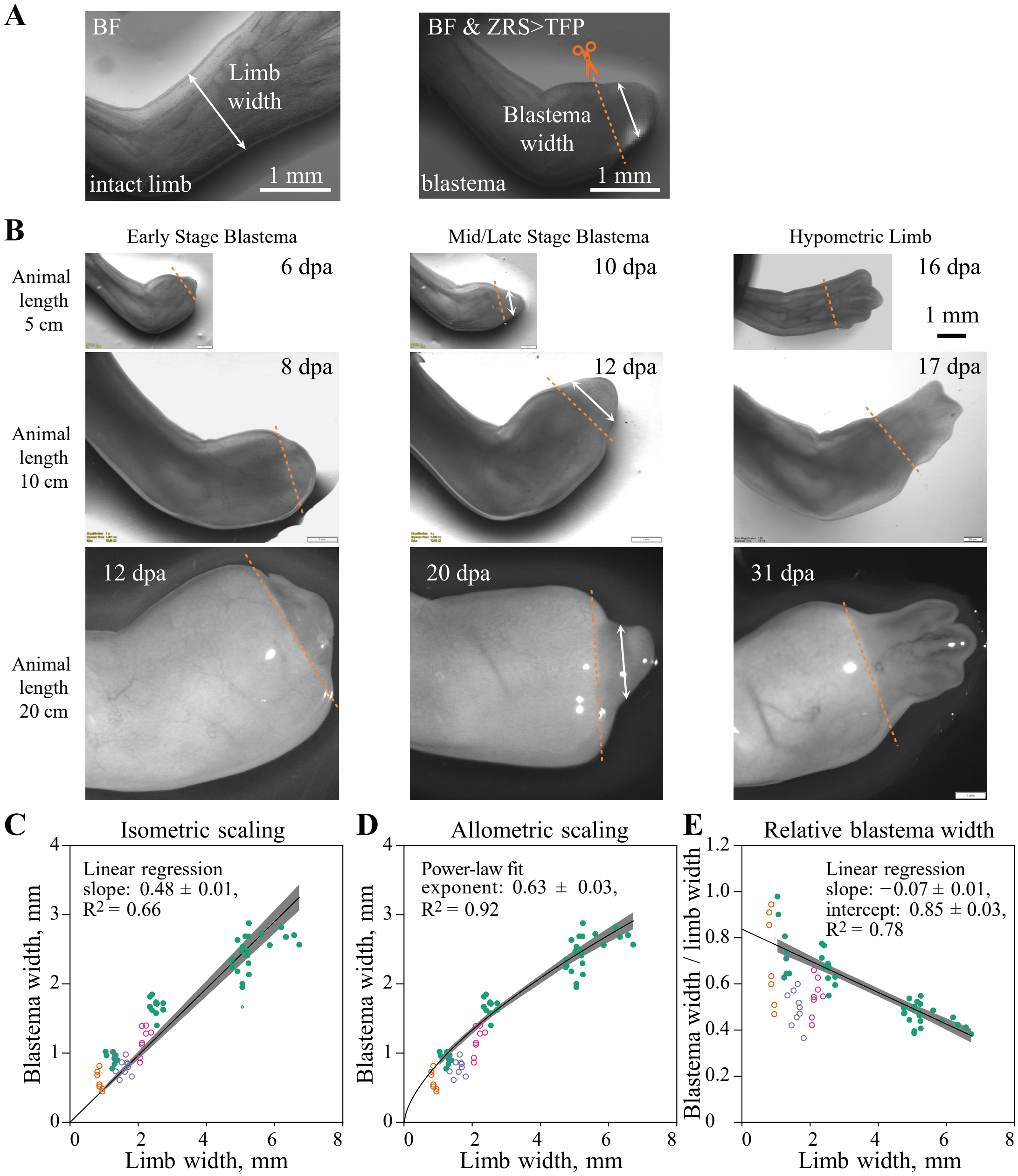


FIG. S2. *Measurements of blastema width.* *A.* Schematic of limb width and blastema width measurements from wide-field images. White double arrows indicate the position of the limb/blastema width measurement, the orange dotted line the amputation plane. Specifically, the position and direction of limb width measurement was chosen to match the later amputation plane. The position of blastema width measurement was chosen as the center of the ZRS>TFP reporter expression, with measurement direction parallel to the amputation plane. *B.* Representative images of blastema in different-sized animals at different time points of regeneration. The amputation plane (i.e., the putative edge of blastema, orange dotted lines) and position of the blastema width measurements (white double arrows) are shown. The discrepancy between hypometric limb width and limb stump width (indicative of initial limb width) is more pronounced in larger animals. *C.* Mid/late-stage blastema width as a function of limb width (green filled symbols), together with proportional fit corresponding to an isometric scaling relation. *D.* Same data as in panel C, but with power-law fit corresponding to an allometric scaling relation. The width of mid/late-stage blastemas scales with exponent $0.63 \pm 0.03$ with intact limb width. Using the fact that the intact limb size scales with animal length with exponent $0.92 \pm 0.01$ (Fig. S3C), we conclude that blastema width scales with animal length with exponent $0.63 * 0.92 = 0.58 \sim 0.6$. *E.* The relative blastema width decreases as a function of limb width (and thus animal size). For comparison, blastema width as shown in Fig. 1C measured for different stages of blastema growth along a line connecting the centers of SHH and FGF8 source regions is shown as open symbols for 3-cm animal at 4-7 dpa (orange), 5-cm animal at 6-10 dpa (blue), 7-cm animal at 8-12 dpa (magenta). Here, we used the linear regression from Fig. S3B to relate animal length and limb width. Note that variability in determining the line connecting the centers of SHH and FGF8 source regions adds to the variability of these blastema width measurements. The corresponding data points were not used for the fits of allometric scaling relations shown. Data: Supplementary Data File Data_S1.xlsx.

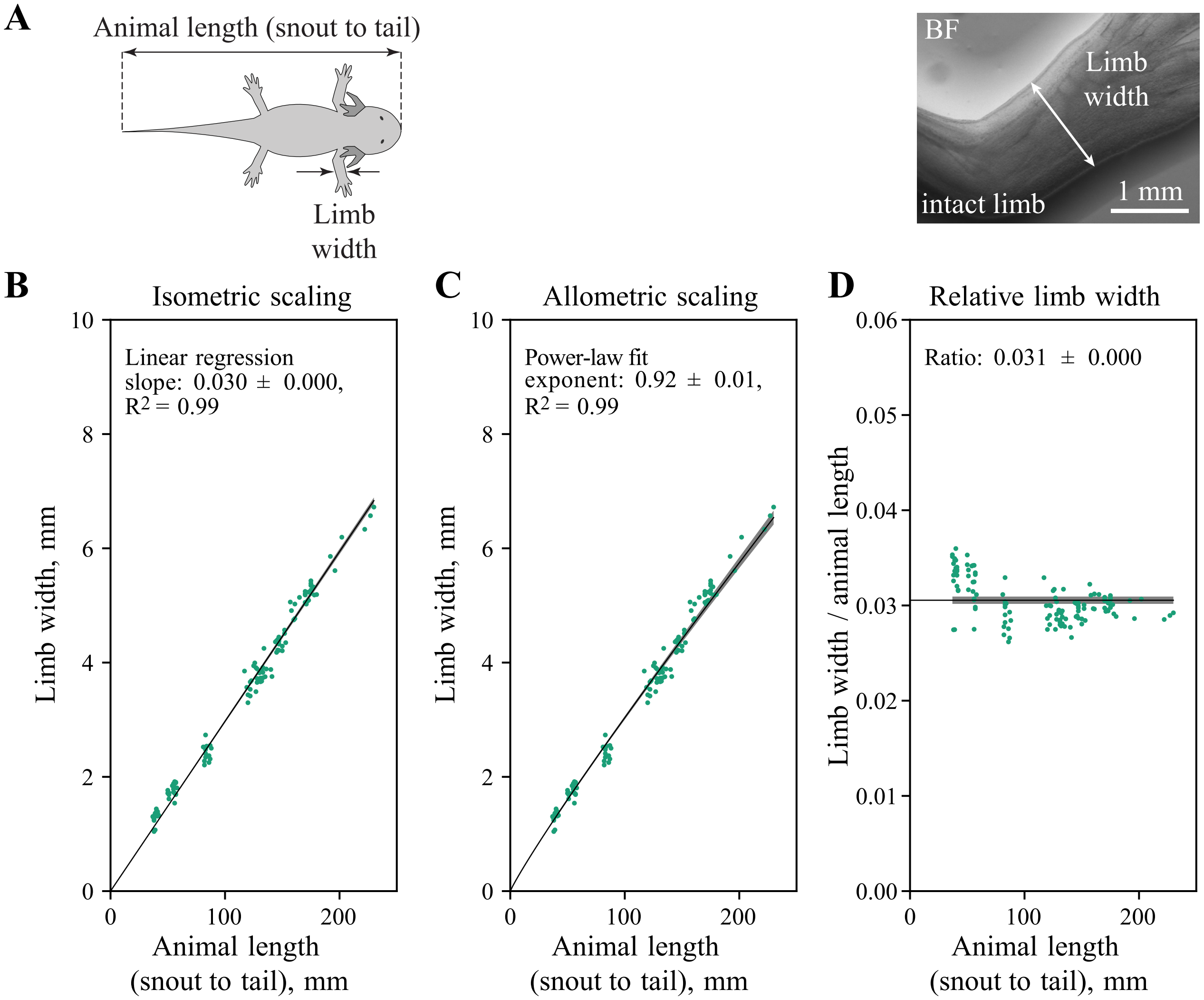


FIG. S3. *Measurements of animal length and limb width. A.* Schematics of animal length measurements (from snout to tail) and limb width measurements from wide-field images; the white double arrow indicates the position of the limb width measurement. *B.* Limb width as a function of animal length, together with proportional fit corresponding to an isometric scaling relation. *C.* Same data as in panel A, but with power-law fit corresponding to an allometric scaling relation. *D.* The relative limb width stays approximately constant as a function of animal length. An analogous analysis using snout-to-cloaca length as alternative measure for animal size gave analogous results. Data: Supplementary Data File Data_S1.xlsx.

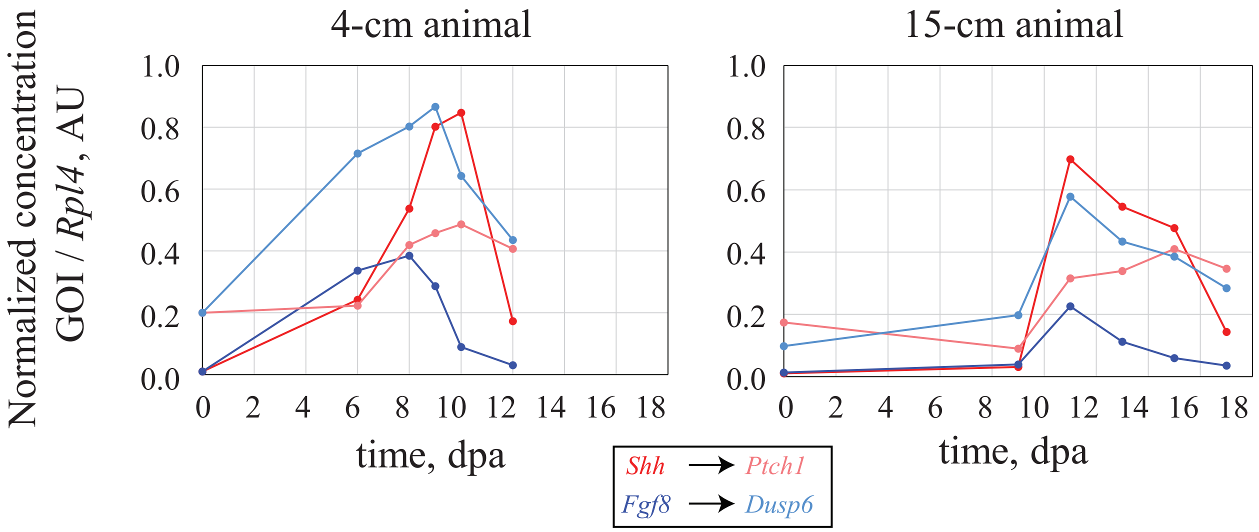


FIG. S4. *Quantitative PCR (qPCR) analysis of Shh, Fgf8, and their downstream targets Ptch1 and Dusp6 during regeneration in animals of two different sizes (snout-to-tail length).* Experimentally measured mRNA values for each Gene of Interest (GOI) were normalized to the housekeeping gene *Rpl4* and presented in arbitrary units. The data represent mean values from two independent experiments. The approximately equal amplitudes of normalized *Shh* and *Fgf8* expression, together with the size-scaling of source regions shown in Fig. 4D, is consistent with the assumption of a size-independent morphogen production rate constant (parameter $\alpha$ in the model).

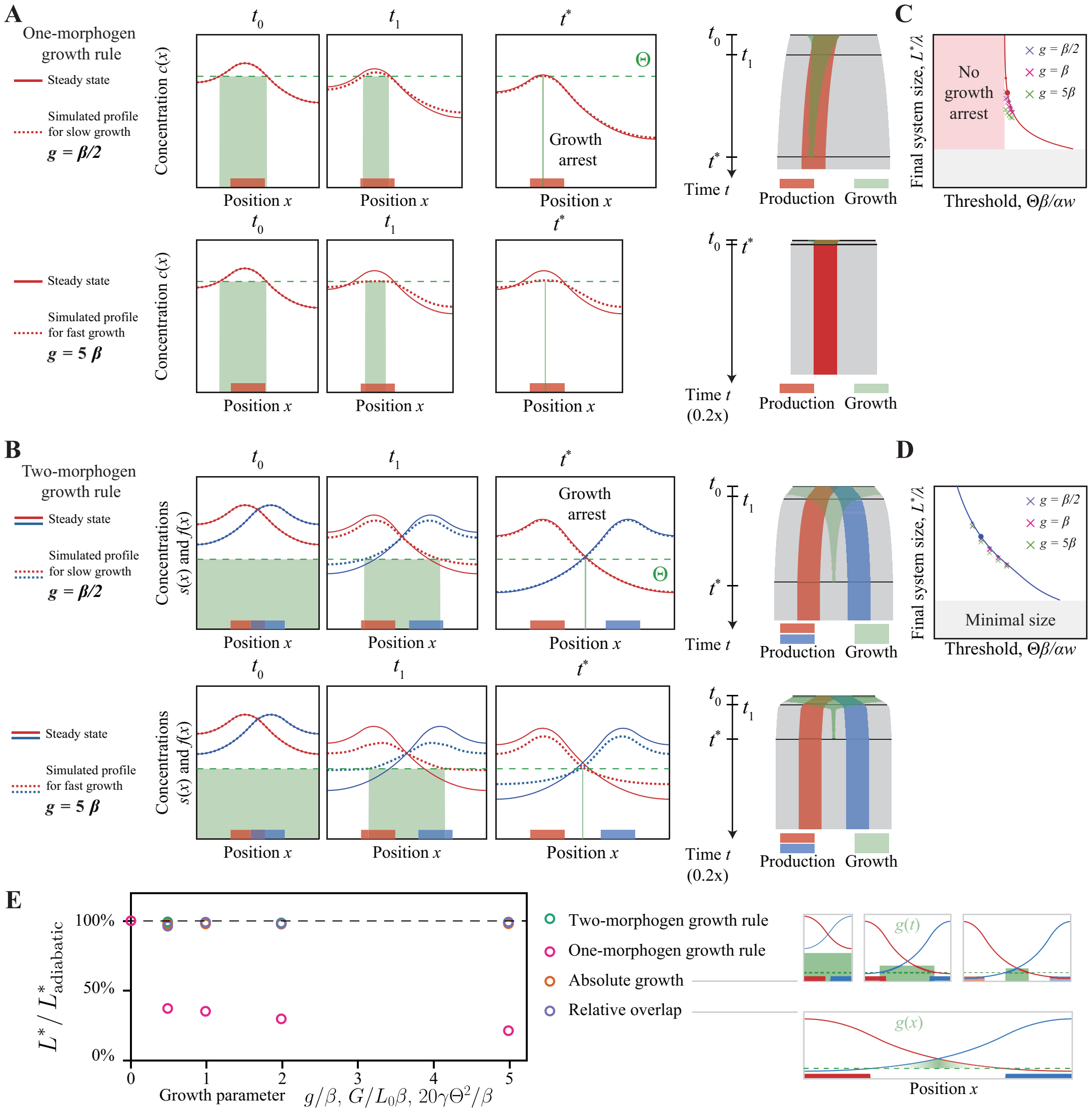


FIG. S5. *Effect of fast growth on final tissue size. A.* Morphogen source region (red region), morphogen gradient (red curve), growth threshold $\Theta$ (dashed green line), and growth zone (green region) at different time points for the one-morphogen growth rule analogous to Fig. 2A, yet for high growth rate $g$, resulting in non-steady-state concentration profiles. Dashed lines show morphogen profiles computed for the case of a finite growth rate $g$ for two values of $g$ (upper and lower panel, respectively), solid lines show morphogen profiles in the limit case of slow growth $g \ll \beta$ as considered in Fig. 2A. Corresponding kymographs of computed tissue growth analogous to Fig. 2A are shown to the right. *B.* Morphogen source region (red region), morphogen gradient (red curve), growth threshold $\Theta$ (dashed green line), and growth zone (green region) at different time points for the two-morphogen growth rule analogous to Fig. 2B, for a finite growth rate $g$. As in panel S5A, dashed lines show morphogen profiles computed for the case of a finite growth rate $g$ for two values of $g$, solid lines show morphogen profiles in the limit case of slow growth as considered in Fig. 2B. Corresponding kymographs of computed tissue growth analogous to Fig. 2B are shown to the right. *C, D.* Final system size $L^*$ normalized by pattern length-scale $\lambda$ as a function of a normalized growth threshold $\Theta\,\beta/\alpha w$ for the one-morphogen growth rule. Also shown are values of final system size for selected growth rates. *E.* Final system size $L^*$ for fast growth, normalized by the system size $L^*_{\text{adiabatic}}$ for the adiabatic limit of slow growth $g \ll \beta$, as a function of the respective growth parameter ($g$, $G$, or $\gamma$) for different growth rules. In addition to the two-morphogen and one-morphogen growth rule introduced in the main text, we investigated two variants of the two-morphogen growth rule with growth rates $g$ that vary either in time or space: *Absolute growth*, where the system grows with a growth rate $g(x) = G/L$ at all positions $x$ where the concentrations $s(x)$ and $f(x)$ of both morphogens exceed the fixed threshold $\Theta$; hence, the absolute change of system size per unit time, normalized by the relative size of the growth zone, is constant. and unit length is constant, as well as a *relative-overlap growth rule*, where the system grows at all positions $x$ where the concentrations $s(x)$ and $f(x)$ exceed the threshold $\Theta$ with a spatially inhomogeneous growth rate $g(x) = \gamma[s(x) - \Theta][f(x) - \Theta]$ that depends on the relative overlap of the two morphogen profiles. We find that the final system size $L^*$ is virtually independent of the rate of growth for the two-morphogen growth rule and its variants, but decreases for faster growth for the one-morphogen growth rule. Parameters: see Table S1, except growth thresholds for one-morphogen growth rule $\Theta = 0.25$, for two-morphogen growth rule $\Theta = 0.1379$.

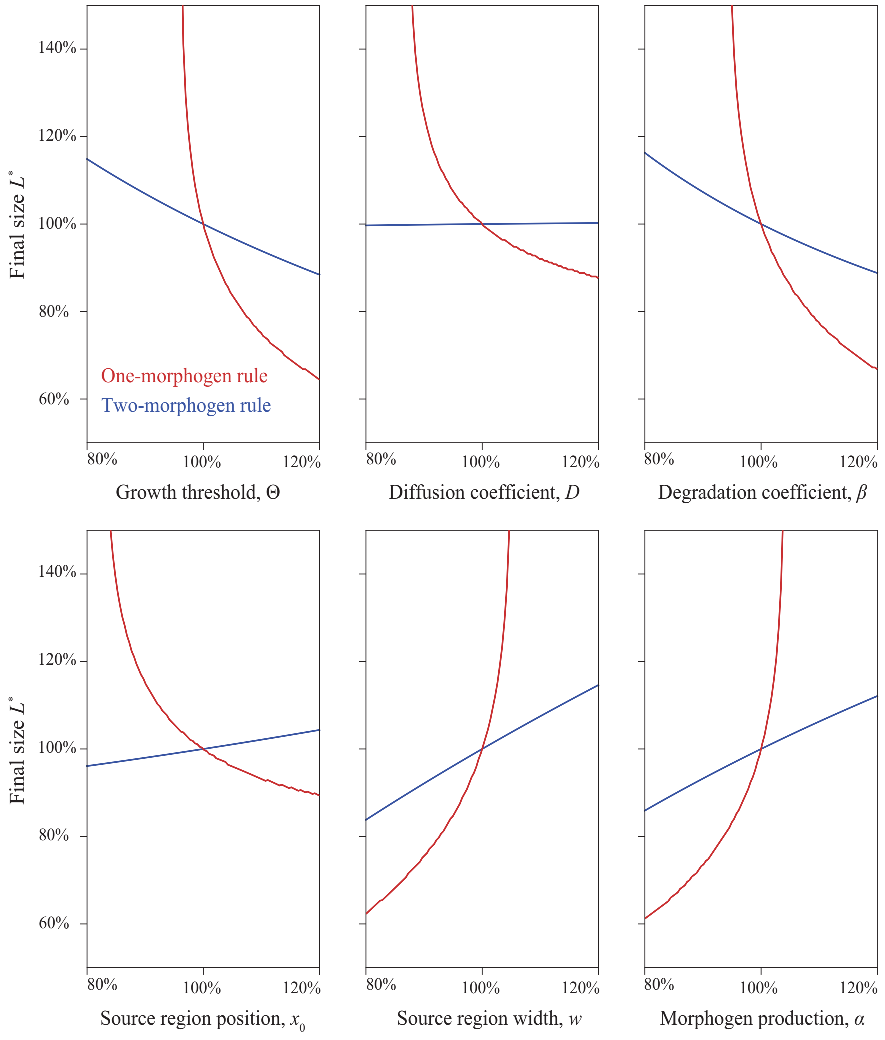


FIG. S6. *Sensitivity of final system size to variations of model parameters.* Plots show the final system size $L^*$ as a function of all model parameters: growth threshold $\Theta$, diffusion coefficient $D$, degradation coefficient $\beta$, source region position $x_0$ and its width $w$, production rate $\alpha$. Red lines correspond to the one-morphogen growth rule and blue to the two-morphogen growth rule. Parameters in Table S1 correspond to 100%; for the growth threshold, we used as reference value $\Theta = 0.25$ for the one-morphogen growth rule, and $\Theta = 0.1379$ for two-morphogen growth rule (same as in Figure S5C-E).

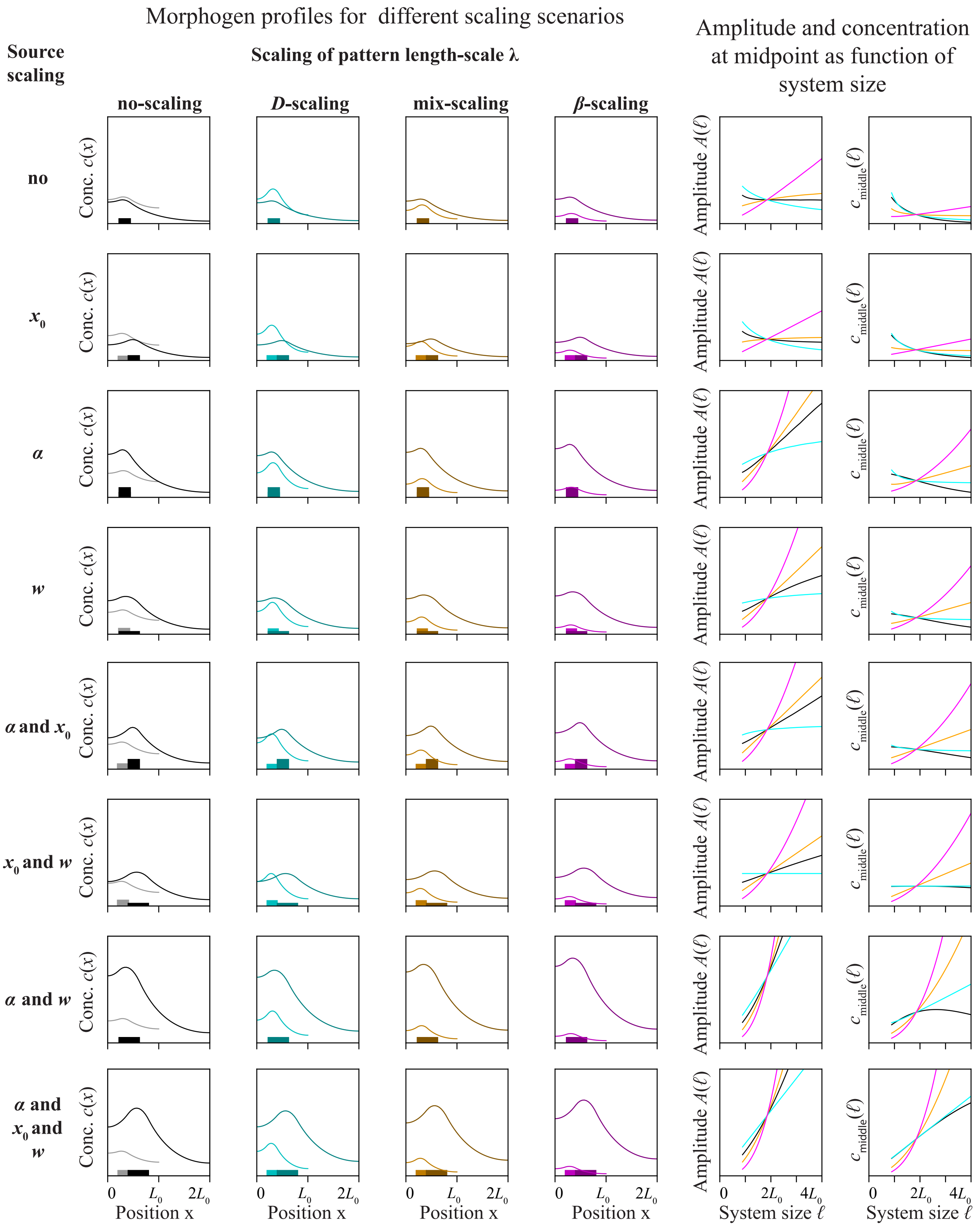


FIG. S7. *Testing all scaling sub-scenarios for 5 model parameters.* The model parameters are: three parameters describing the source region: source width $w$, source position $x_0$, production rate $\alpha$; and two parameters defining the pattern length-scale $\lambda = \sqrt{D/\beta}$, which can scale according to either $D$-scaling ($D \sim \ell^2$), mixed-scaling ($D \sim \ell$ and $\beta \sim 1/\ell$), or $\beta$-scaling ($\beta \sim 1/\ell^2$). The left side of the figure shows the overlaid morphogen profiles for two system sizes $\ell_0$ and $2\ell_0$, (analogous to Fig. 2A) for each scaling sub-scenarios corresponding to the different combinations of source-parameter scaling (rows) and pattern length-scale-scaling (columns). The right side shows the amplitude and the concentration at the midpoint of morphogen profiles as a function of system size $\ell$ for different scaling sub-scenarios, colored analogously to the left panel. To enable growth arrest in the case of the one-morphogen growth rule, the morphogen gradient amplitude must decay with $\ell$; in the case of the two-morphogen growth rule, the morphogen concentration at the midpoint must decay with $\ell$. Parameters: see Table S1.

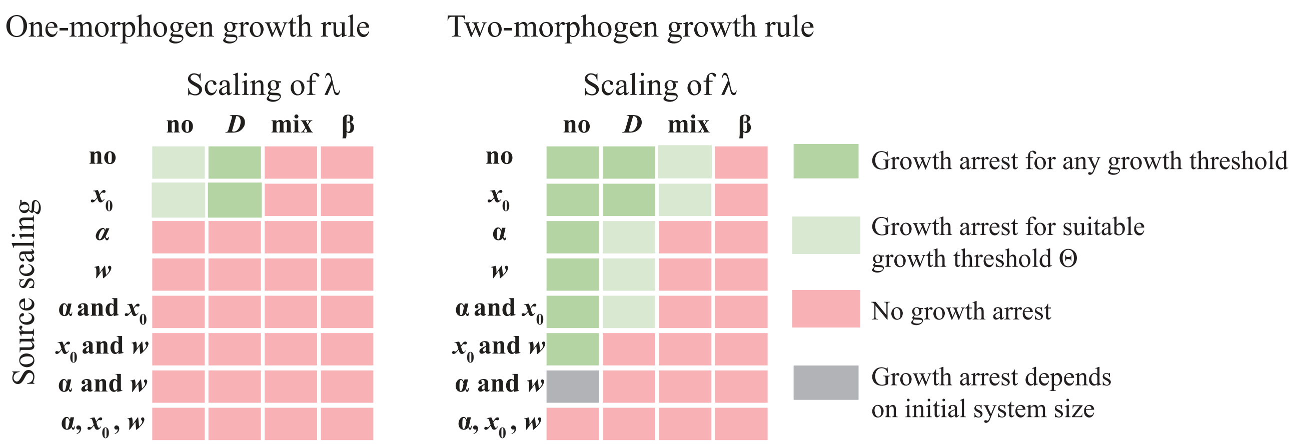


FIG. S8. *Graphical summary of growth arrest for all scaling sub-scenarios for scaling of 5 model parameters for the scenario of dynamic scaling with blastema size.* Rows correspond to different sub-scenarios of source parameters (position $x_0$, width $w$, production rate $\alpha$) that may scale with blastema size $L(t)$; columns correspond to different sub-scenarios of morphogen gradient parameters that may scale with $L(t)$, analogous to Fig. S7. For each sub-scenario of parameter scaling, it is indicated whether growth arrests for all choices of the growth threshold $\Theta$ (green), growth arrests if the growth threshold $\Theta$ exceeds a critical value $\Theta_c$ (light green), growth never arrests (red), or growth arrests only if the initial system size is smaller than a critical system size (gray). Results correspond to source position $x_0 > 0$ and source size $w > 0$ as in Fig. S7; results for the limit case of a point source located at the system's boundary are very similiar, see Table S2.

**A**

**Allometric scaling exponent ν**

Scaling of pattern length-scale λ

| Source scaling | no scaling | *D*-scaling | mix-scaling | *β*-scaling |
|---|---|---|---|---|
| no | - | 0.1 | 0.5 - 1.0 | 1.4 - 1.9 |
| $x_0$ | 0.4 - 0.8 | 0.2 - 0.8 (B) | 0.8 - 1.0 | 1.2 - 3.1 |
| $w$ | 0.6 - 1.0 (A) | 0.5 - 1.0 | 1.0 - 2.1 | 1.3 - 3.6 |
| $x_0$ and $w$ | 0.9 -1.0 | 1.0 (I) | 1.1 - 2.0 | 1.3 - 3.1 |

Allometric $L^* \sim L_a^\nu$ — Isometric $L^* \sim L_a$

**B**

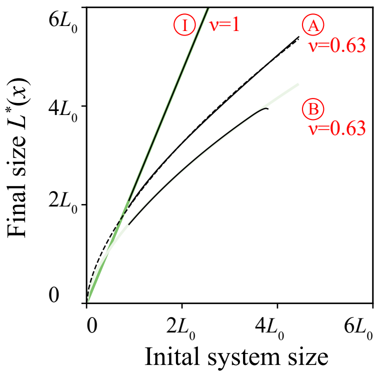


FIG. S9. *Graphical summary of allometric scaling relations for static scaling with animal size.* *A*. Predicted scaling exponents $\nu$ of allometric scaling relations $L^* \sim L_a^\nu$ between final system size $L^*$ and initial system size (assumed proportional to animal size $L_a$) for different sub-scenarios of static scaling with animal size for the two-morphogen growth rule. We systematically tested combinations of different sub-scenarios for the scaling of source size parameters with animal size (position $x_0$, width $w$; rows) and the scaling of the pattern length-scale $\lambda$ with animal size $L_a$ (columns), and report the range of exponents $\nu$ obtained by fitting a power law $L^* \sim L_a^\nu$ to solutions of Eq. S29 for different values for $\lambda$, $w$, $x_0$ (specified below). Only fits with a determination coefficient $R^2 > 0.95$ were included. *B*. Final system size $L^*$ as a function of initial system size for three selected scaling sub-scenarios (highlighted in panel A): I: scaling of source position and width combined with $D$-scaling of pattern length-scale $\lambda$, resulting in proportional growth, A: scaling of source width without scaling of $\lambda$, and B: scaling of source position combined with $D$-scaling. $R^2$ was 1.0 in all three cases; the allometric scaling exponent $\nu = 0.63$ for A and B, which is close to the allometric scaling exponent observed for late-bud-stage blastemas [8] (see also Fig. S2). (Note that a more accurate comparison would refer to the ratio of allometric scaling exponents at the beginning and end of the first phase of blastema outgrowth, corresponding to the relative growth during this first growth phase. From the scaling exponents 0.61 and 0.43 for late-bud-stage blastemas and blastemas during subsequent digit cartilage formation several days later, respectively [8], we conclude that this ratio is likely smaller than 0.70=0.43/0.61.) Parameters: see Table S1, with growth threshold $\Theta = 0.15$; in panel A, $\lambda/L_a$ was varied in the range 0.01-0.9, $w/L_a = x_0/L_a$ was varied in the range 0-0.5; in panel B, $\lambda/L_a = 0.18$ and $w/L_a = x_0/L_a = 0.48$.

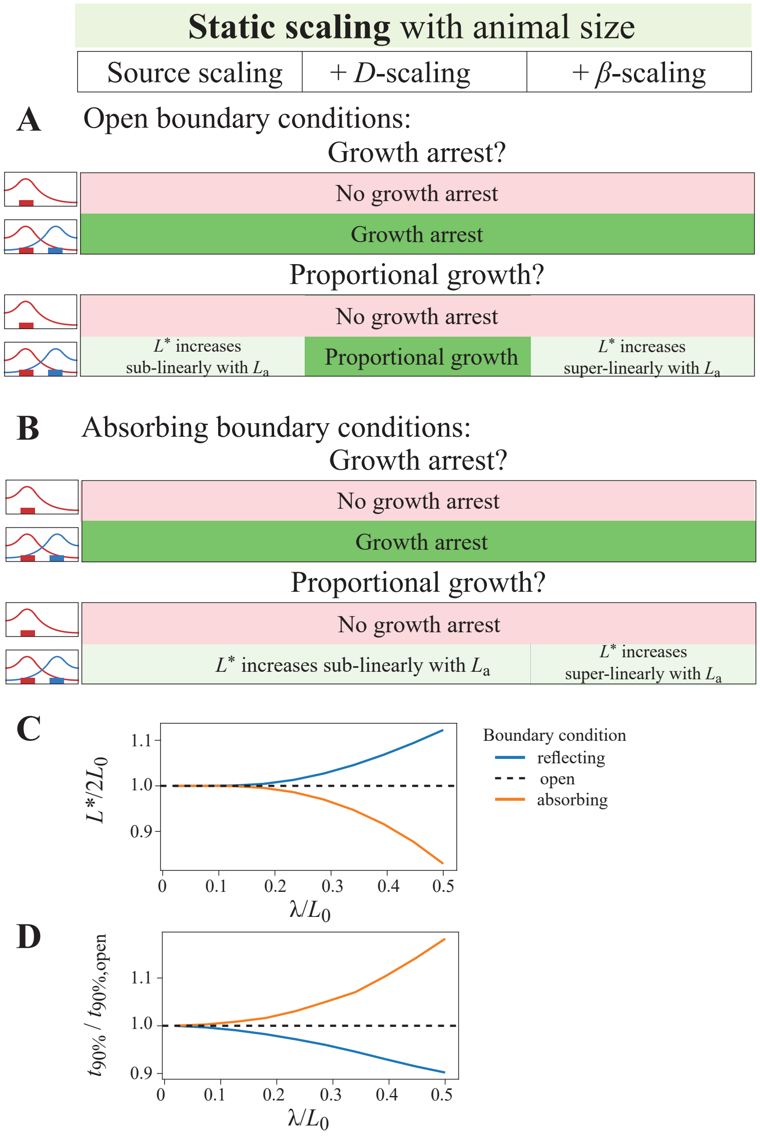


FIG. S10. *Growth arrest and proportional growth for static scaling scenario for different boundary conditions. A, B.* Analysis analogous to Fig. 3D, yet assuming either (A) open boundary conditions (by placing the finite system of size $L$ inside a larger system), or (B) absorbing boundary conditions (by imposing the boundary conditions $s(x{=}0) = f(x{=}0) = s(x{=}L) = f(x{=}L) = 0$) instead of the reflecting boundary conditions used for Fig. 3D). Note that for the dynamic scaling scenario, results are identical to those of Fig. 3C when using either open or absorbing boundary conditions. *C.* Final system size $L^*$ as a function of pattern length-scale $\lambda$ for different boundary conditions assuming the two-morphogen growth rule. To ensure comparability for different values of $\lambda$, the growth threshold $\Theta$ was adapted for each $\lambda$, such that the system grows exactly to twice its initial size $L_0$ for the reference case of open boundary conditions (i.e., $L^* = 2L_0$ in this case). *D.* Typical growth time $t_{90\%}$ as a function of $\lambda/L_0$ for the computations shown in panel C. Here, $t_{90\%}$ denotes the time until which 90% of the total increase in system size has been completed (i.e., $L(t_{90\%}) = L_0 + 0.9(L^* - L_0)$). The case of open boundary conditions is again used as reference case, with typical growth times $t_{90\%}$ normalized by the corresponding value $t_{90\%,\text{open}}$ for open boundary conditions. Parameters except $\Theta$: see Table S1.

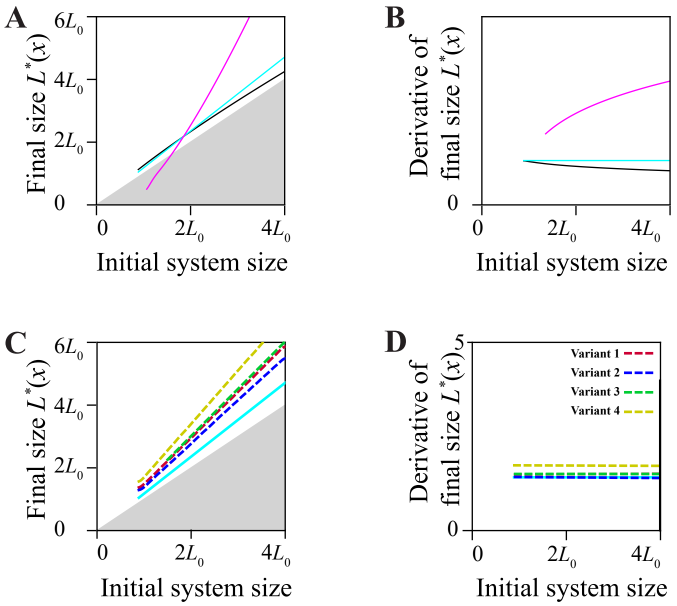


E

| Parameter | Variant 1 | Variant 2 | Variant 3 | Variant 4 |
|---|---|---|---|---|
| $w$ for SHH | dynamic | dynamic | dynamic | dynamic |
| $x_0$ for SHH | dynamic | dynamic | dynamic | dynamic |
| $\lambda$ for SHH | dynamic | static | static | dynamic |
| $w$ for FGF8 | static | static | static | static |
| $x_0$ for FGF8 | static | static | dynamic | dynamic |
| $\lambda$ for FGF8 | static | dynamic | dynamic | dynamic |

| | Growth arrest? |
|---|---|
| Two-morphogen | for $D$-scaling |

| | Proportional growth? |
|---|---|
| Two-morphogen | for $D$-scaling |

FIG. S11. *Proportional growth for mixed scaling scenarios.* For reference, panels A and B address scenarios of only static scaling of morphogen gradient parameters as shown in Fig. 2 in the main text, while panels C, D and E address variants of mixed scaling in which some parameters display dynamic scaling and others exhibit static scaling. *A.* Final system size as a function of initial system size (assumed proportional to $L_a$) for three gradient scaling sub-scenarios as in Fig. 2: source scaling (black), $+D$-scaling (cyan), $+\beta$-scaling (magenta). *B.* Numerical derivative of the final system size with respect to the initial system size for the data shown in Panel A, to discriminate between sub-linear growth (decreasing), proportional growth (constant), and super-linear growth (increasing). *C.* Final system size as a function of initial system size for mixed scaling scenarios combining static and dynamic scaling (listed in panel E, assuming $D$-scaling; dashed), as well as the result for static scaling with $D$-scaling from panel A for comparison (cyan). *D.* Numerical derivative of final system size with respect to initial system size for the data shown in Panel C, revealing a constant slope indicative of perfect proportional growth as implied by analytical theory, see section ‘Mixed scaling variants for $D$-scaling’. *E.* Variants of mixed scaling scenarios, where some morphogen gradient parameters exhibit dynamic scaling, while other parameters exhibit static scaling. For each variant, our model predicts growth arrest and proportional growth if the two-morphogen growth rule and $D$-scaling is assumed. Parameters: see Table S1, growth threshold $\Theta = 0.15$.

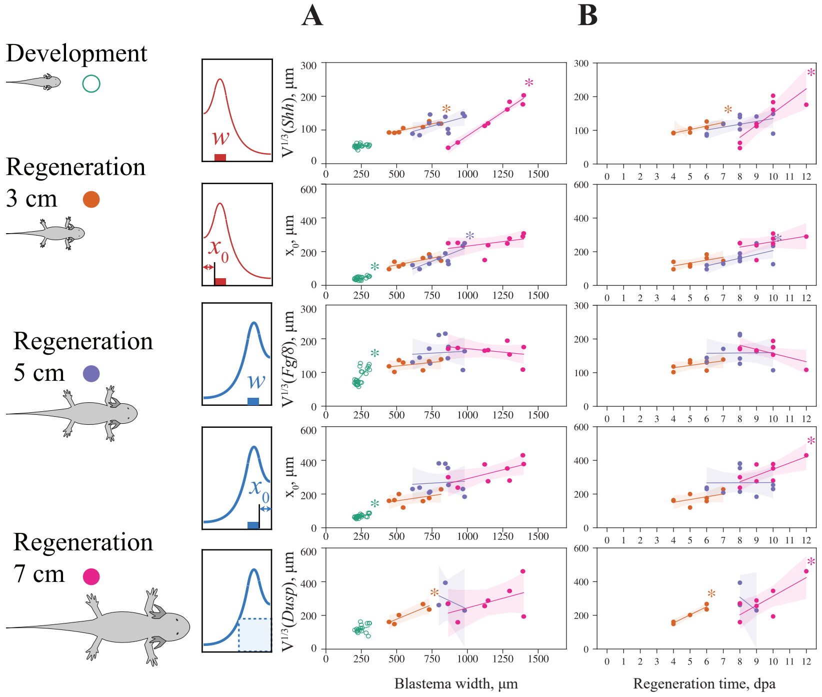


FIG. S12. *Additional fit variant for Fig. 4D. A.* Same data as in Fig. 4D for experimentally measured characteristics of SHH and FGF8 source regions and signaling range, yet with separate linear regressions for each animal-size group. If a morphogen parameter displays dynamic scaling, regressions should have a significant positive slope, while for static scaling, the individual regressions should have a slope that is not statistically different from zero. *B.* Same data, yet with separate linear regressions for each animal-size group plotted against regeneration time. Again, if a morphogen parameter displays dynamic scaling, regressions should have a significant positive slope, while for static scaling, the individual regressions should have a slope that is not statistically different from zero. Stars indicate that the fitted slope from linear regression is statistically significantly different from zero ($p < 5\%$).

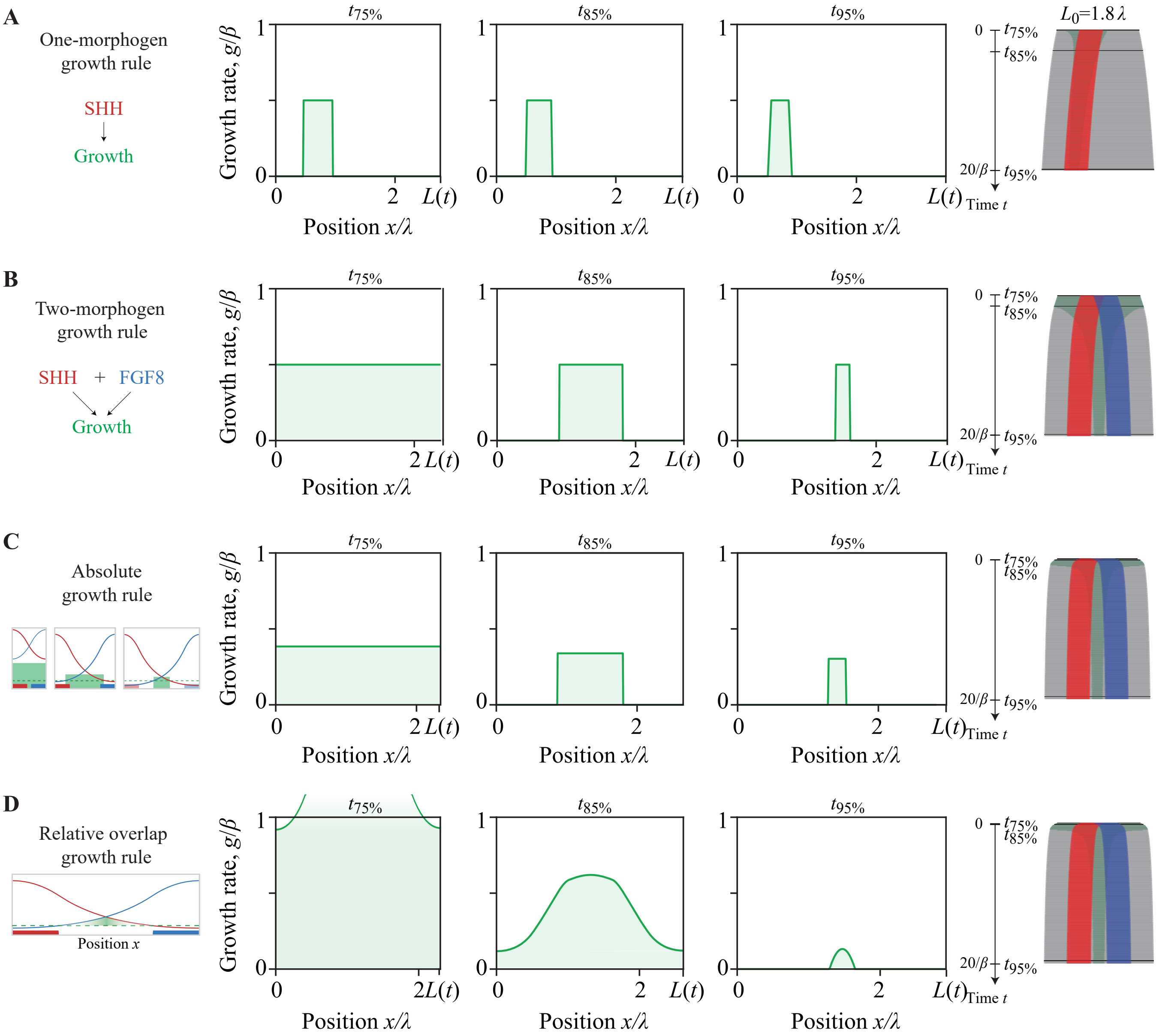


FIG. S13. *Spatial growth profiles for different growth rules. A.* Spatial profile of growth rate $g(x,t)$ (normalized by degradation rate $\beta$) as a function of position $x$ (normalized by pattern length-scale $\lambda$) for the *one-morphogen growth rule* as introduced in Fig. 2A at three subsequent time points $t_{75\%}$, $t_{85\%}$, $t_{95\%}$, at which the system has grown 75%, 85%, 95% of the total increase in system size $L^* - L_0$, respectively (e.g., $L(t_{75\%}) - L_0 = 75\%\,(L^* - L_0)$). The corresponding kymograph of simulated tissue growth (analogous to Fig. 2A) is shown to the right. *B.* Analogous to panel A, but for the *two-morphogen growth rule* as introduced in Fig. 2B. *C.* Analogous to panel A, but for the *absolute growth rule* as introduced in Fig. S5E. Note the decrease of the growth rate inside the growth zone as a function of time. *D.* Analogous to panel A, but for the *relative overlap growth rule* as introduced in Fig. S5E. Parameters: see Table S1; $g/\beta = 0.5$ (panels A and B), $G = 0.5\,L_0\beta$ (panel C), $\gamma = 0.5\,\beta/20\Theta^2$ (panel D).

| **Parameter** | **Value** |
|---|---|
| Diffusion coefficient $D$ | $D_0 = 1$ |
| Degradation rate $\beta$ | $\beta_0 = 1$ |
| Pattern length-scale $\lambda$ | $\lambda_0 = \sqrt{D_0/\beta_0} = 1$ (Fig. 2, Fig. 3A-left) |
| Production rate $\alpha$ | $\alpha_0 = 1$ |
| Growth rate $g$ | $0.5\,\beta_0$ |
| Source position $x_0$ | 0.4 (Fig. 2) |
| Source width $w$ | 0.4 (Fig. 2) |
| Source position $x_0(L)$ for *dynamic* scaling | $x_0 = 0.13\,L(t)$ (Fig. 3A), $x_0 = 0.2\,L(t)$, (Fig. S7) |
| Source width $w(L)$ for *dynamic* scaling | $w = 0.13\,L(t)$ (Fig. 3A), $w = 0.2\,L(t)$ (Fig. S7) |
| Source position $x_0(L_0)$ for *static* scaling | $x_0 = 0.2\,L_0$ (Fig. 3B, Fig. S7) |
| Source width $w(L_0)$ for *static* scaling | $w = 0.2\,L_0$ (Fig. 3B, Fig. S7) |
| Pattern length-scale for *dynamic* source scaling | $\lambda = \lambda_0$ (Fig. 3A-left) |
| Diffusion coefficient $D(L)$ for *dynamic* $D$-scaling | $D = [0.3\,L(t)]^2\beta_0$ (Fig. 3A-middle, Fig. S7) |
| Degradation rate $\beta(L)$ for *dynamic* $\beta$-scaling | $\beta = D_0/[0.3\,L(t)]^2$ (Fig. 3A-right, Fig. S7) |
| Pattern length-scale $\lambda$ for *static* source scaling | $\lambda = 1.8$ (Fig. 3B-left, Fig. 3B-upper row) |
| Diffusion coefficient $D(L_0)$ for *static* $D$-scaling | $D = L_0^2\beta_0$ (Fig. 3B-middle)<br>$D = [0.3\,L_0]^2\beta_0$ (Fig. S7) |
| Degradation rate $\beta(L_0)$ for *static* $\beta$-scaling | $\beta = D/L_0^2$ with $D = 1.8^2\beta_0$ (Fig. 3B-right)<br>$\beta = D_0/[0.3\,L_0]^2$ (Fig. S7) |
| Initial system size $L_0$ | 1.8<br>Fig. 3B-lower row: 3.6 |
| Growth threshold $\Theta$ | Fig. 2A: 0.25<br>Fig. 2B: 0.14<br>Fig. 3B: 0.1 |
| Feedback competent region size $w_{\text{comp.}}$ | $0.8\,\lambda_0$ (Fig. 5) |

TABLE S1. *Dimensionless parameter values used in numerical computations.* The morphogen parameters (diffusion coefficient $D$, degradation rate $\beta$, and production rate $\alpha$) were set equal to 1; different values for these parameters correspond to trivial re-scaling of length units, time units, concentration units, and do not change any of our conclusions, see section 'Re-scaling of units'. In Fig. 2, lengths are reported relative to the characteristic length scale $\lambda_0 = 1$, concentrations are reported relative to the characteristic concentration $\alpha_0 w_0/\beta_0$. The growth rate $g$ was set to $g = 0.5\beta_0$ for most figures; other values of $g$ are tested in Fig. S5. The source parameters (source position $x_0$, source width $w$) and the proportionality coefficient assumed in morphogen gradient scaling were chosen to mimic the experimentally found values of these parameters (Fig. 4), normalized by the experimental estimate of FGF8 pattern length-scale.

| | *Dynamic* scaling with source scaling $w \sim L$ | | | |
|---|---|---|---|---|
| | No scaling<br>$\lambda = \text{const}$ | $D$-scaling<br>$D \sim L^2$ | Mixed-scaling<br>$D \sim L; \beta \sim 1/L$ | $\beta$-scaling<br>$\beta \sim 1/L^2$ |
| Characteristic concentration, $\Theta_c = aw/(D\beta)^{1/2}$ | $\sim L$ | const | $\sim L$ | $\sim L^2$ |
| Amplitude, $A = \max_{x\in[0,L]} c^*(x) = \Theta_c \coth L/\lambda$ | $\sim L \coth(CL)$ | const | $\sim L$ | $\sim L^2$ |
| Concentration at $x = L/2$, $c^*(L/2) = \Theta_c/2 \operatorname{cosech} L/2\lambda$ | $\sim L \operatorname{cosech}(CL)$ | const | $\sim L$ | $\sim L^2$ |
| Final system size, one-morphogen growth rule: | – | – | – | – |
| Final system size, two-morphogen growth rule: | const | – | – | – |

| | *Dynamic* scaling without source scaling $w$=const | | | |
|---|---|---|---|---|
| | No scaling | $D$-scaling | Mixed-scaling | $\beta$-scaling |
| Characteristic concentration $\Theta_c$ | const | $\sim L^{-1}$ | const | $\sim L$ |
| Amplitude $A$ | $\sim \coth(CL)$ | $\sim L^{-1}$ | const | $\sim L$ |
| Concentration at midpoint $c(L/2)$ | $\sim \operatorname{cosech}(CL)$ | $\sim L^{-1}$ | const | $\sim L$ |
| Final system size, one-morphogen growth rule: | const if $\Theta > \Theta_c$ | const | – | – |
| Final system size, two-morphogen growth rule: | const | const | – | – |

| | *Static* scaling with source scaling $w \sim L_0$ | | | |
|---|---|---|---|---|
| | No scaling<br>$\lambda = \text{const}$ | $D$-scaling<br>$D \sim L_0^2$ | Mixed-scaling<br>$D \sim L_0; \beta \sim 1/L_0$ | $\beta$-scaling<br>$\beta \sim 1/L_0^2$ |
| Characteristic concentration $\Theta_c$ | $\sim L_0$ | const | $\sim L_0$ | $\sim L_0^2$ |
| Amplitude $A$ | $\sim L_0 \coth(CL)$ | $\sim \coth(CL/L_0)$ | $\sim L_0 \coth(CL/L_0)$ | $\sim L_0^2 \coth(CL/L_0)$ |
| Concentration at midpoint $c(L/2)$ | $\sim L_0 \operatorname{cosech}(CL)$ | $\sim \operatorname{cosech}(CL/L_0)$ | $\sim L_0 \operatorname{cosech}(CL/L_0)$ | $\sim L_0^2 \operatorname{cosech}(CL/L_0$ |
| Final system size, one-morphogen growth rule:<br>$L^* = \lambda \operatorname{arcoth}(\Theta/\Theta_c)$ | $\sim \operatorname{arcoth}(C/L_0)$ | $\sim L_0$ for $\Theta > \Theta_c$ | $\sim L_0 \operatorname{arcoth}(C/L_0)$ | $\sim L_0 \operatorname{arcoth}(C/L_0^2)$ |
| Final system size, two-morphogen growth rule:<br>$L^* = 2\lambda \operatorname{arsinh}(\Theta_c/2\Theta)$ | $\sim \operatorname{arsinh}(CL_0)$ | $\sim L_0$ | $\sim L_0 \operatorname{arsinh}(CL_0)$ | $\sim L_0 \operatorname{arsinh}(CL_0^2)$ |

| | *Static* scaling without source scaling $w$=const | | | |
|---|---|---|---|---|
| | No scaling | $D$-scaling | Mixed-scaling | $\beta$-scaling |
| Characteristic concentration $\Theta_c$ | const | $\sim L_0^{-1}$ | const | $\sim L_0$ |
| Amplitude $A$ | $\sim \coth(CL)$ | $\sim L_0^{-1} \coth(CL/L_0)$ | $\sim \coth(CL/L_0)$ | $\sim L_0 \coth(CL/L_0)$ |
| Concentration at midpoint $c(L/2)$ | $\sim \operatorname{cosech}(CL)$ | $\sim L_0^{-1} \operatorname{cosech}(CL/L_0)$ | $\sim \operatorname{cosech}(CL/L_0)$ | $\sim L_0 \operatorname{cosech}(CL/L_0$ |
| Final system size, one-morphogen growth rule: | const if $\Theta > \Theta_c$ | $\sim L_0 \operatorname{arcoth}(CL_0)$ | $\sim L_0$ | $\sim L_0 \operatorname{arcoth}(C/L_0)$ |
| Final system size, two-morphogen growth rule: | const | $\sim L_0 \operatorname{arsinh}(C/L_0)$ | $\sim L_0$ | $\sim L_0 \operatorname{arsinh}(CL_0)$ |

TABLE S2. *Summary of analytical results.* In the limit case of a point source region, $w \to 0$, located at the left system boundary, $x_0 = 0$, the steady-state solution of the diffusion equation Eq. (1) is given by the analytical formula Eq. (S20). This table summaries how for Eq. (S20) the amplitude of the morphogen gradient profile and its concentration at the midpoint of the system depend on system size $L$ for different sub-scenarios of dynamic scaling. Additionally, it is stated how the final system size $L^*$ depends on the initial system size $L_0$ for different sub-scenarios of static scaling. Here, $C$ denotes size-independent constants.

| | Dynamic scaling with blastema size? *Null hypothesis:* static scaling | | Static scaling with animal size? *Null hypothesis:* dynamic scaling | |
|---|---|---|---|---|
| Morphogen gradient parameter | Partial correl. analysis $\eta^2$ | $p$ | Partial correl. analysis $\eta^2$ | $p$ |
| $w$ for SHH | 0.58 | <0.1% | 0.19 | n.s. |
| $x_0$ for SHH | 0.54 | <0.1% | <0.01 | n.s. |
| $w$ for FGF8 | <0.01 | n.s. | 0.15 | 0.3% |
| $x_0$ for FGF8 | 0.14 | 0.5% | 0.19 | 0.1% |
| $V^{1/3}$ for *Dusp6* | 0.17 | 0.9% | <0.01 | n.s. |
| $L_a$ + noise | [-0.12,0.12] | n.s. | [0.25,0.48] | [<0.1%,5%] |
| $L$ + noise | [0.24,0.47] | [<0.1%,6%] | [-0.13,0.14] | n.s. |

TABLE S3. *Partial correlation analysis for morphogen gradient parameters against animal and blastema size.* Shown are the results of a partial correlation analysis for morphogen gradient parameters as reported in Fig. 4D, which allows testing for conditional statistical independence. To test for *dynamic scaling*, we aim to refute a null hypothesis of static scaling. We performed a partial correlation analysis against blastema size, after factoring out animal size. If the null hypothesis were true, no correlation should remain. Here, $\eta^2$ denotes the explained residual variance of a morphogen gradient parameter $X$ due to the influence of blastema size $Y = L$ after factoring out the influence of animal size $Z = L_a$ according to Eq. S8, while $p$ denotes the significance level of a positive linear correlation between blastema size and a morphogen parameter after factoring out animal size, computed using Eq. S11. A non-significant $p$-value is consistent with static scaling of this parameter, while a high $\eta^2$ and statistically significant $p$ is indicative of *dynamic scaling.* To test for *static scaling*, we aim to refute a null hypothesis of dynamic scaling. We analogously performed a partial correlation analysis against animal size, after factoring out blastema size. Here, $\eta^2$ denotes the explained residual variance of a morphogen gradient parameter due to the influence of animal size $Y = L_a$ after factoring out the influence of blastema size $Z = L$, and $p$ again the significance level of the corresponding correlation. A non-significant $p$-value is consistent with dynamic scaling of this parameter. For each morphogen parameter, $n = 48$ samples were analyzed, except $n = 33$ for *Dusp6*. Animal size corresponds to snout-to-tail animal length; snout-to-cloaca animal length gave very similar results. For comparison, we applied the same test also to synthetic data comprising $n = 48$ samples with the same animal and blastema sizes as measured in the experiments, and a hypothetical parameter $X$ given by either animal size $L_a$ or blastema size $L$ multiplied by independent, normally distributed multiplicative noise factors with mean 1 and variance 0.25. Intervals for $\eta^2$ corresponds to the $[25\%, 75\%]$-inter quartile range.

### I. SUPPLEMENTARY METHODS

#### A. Numerical methods

As mentioned in the Methods section of the main text, in the presence of tissue growth with local growth rate $g(x,t)$, Eq. 1 changes to Eq. 4

$$\partial_t c(x,t) = D\,\partial_x^2 c(x,t) - v(x,t)\partial_x c(x,t) - [\beta + g(x,t)]c(x,t) + \alpha\,\chi_w(t)(x) \quad , \tag{S1}$$

where $\partial_x v = g$. Here, the growth-induced "dilution" term $-gc$ accounts for the fact that in a growing tissue the same amount of morphogen will be spread in an increasingly larger system, which decreases the local concentration $c$ at a rate equal to the growth rate $g$ [21, 46]. To derive this term, note that Eq. S1 can be equivalently written as a continuity equation $\partial_t c = -\partial_x J - \beta c + \alpha\chi_w$, where $J = -D\partial_x c + vc$ is the combined current of morphogen molecules due to both diffusion with diffusion coefficient $D$ and growth-induced advection with velocity $v$. By the product rule, $\partial_x(vc) = v(\partial_x c) + (\partial_x v)c = v\,\partial_x c + gc$ because $\partial_x v = g$, which naturally gives rise to the dilution term $-gc$ in Eq. S1.

This equation is numerically solved using a modified Euler scheme in a Lagrangian frame. While Eulerian coordinates describe fixed coordinate positions that are stationary with respect to a fixed laboratory frame, Lagrangian coordinates describe moving coordinates that track a particular point of a flowing material such as a growing tissue and thus move with its flow. Specifically, we consider a non-uniform spatial grid with grid points $x_i(t)$ in one space dimension, whose bin sizes $\Delta x_i(t_n) = x_{i+1}(t_n) - x_i(t_n)$ change in time according to

$$\Delta x_i(t_{n+1}) = [1 + g(x_i,t_n)\Delta t]\Delta x_i(t_n) \quad . \tag{S2}$$

The discretization of the Laplacian $\partial_x^2 c$ for this non-uniform grid reads

$$\partial_x^2 c(x_i,t_n) \approx \frac{2}{x_{i+1}(t_n) - x_{i-1}(t_n)}\left(\frac{c(x_{i+1},t_n) - c(x_i,t_n)}{x_{i+1}(t_n) - x_i(t_n)} - \frac{c(x_i,t_n) - c(x_{i-1},t_n)}{x_i(t_n) - x_{i-1}(t_n)}\right) \quad , \tag{S3}$$

where $c(x_i,t_n)$ is the concentration at position $x_i$ and time $t_n$ (more precisely, the average concentration in a spatial bin with center $x_i$), and $x_i$ specifically denotes in this case the center of the spatial bin with the varying size. Note that the above Laplacian for the non-uniform grid reduces to the standard second-central-difference formula $c(x_{i+1},t_n) - 2c(x_i,t_n) + c(x_{i-1},t_n)/\Delta x^2$ if the grid spacing is uniform $x_{i+1}(t_n) - x_i(t_n) = x_i(t_n) - x_{i-1}(t_n) = \Delta x$.

For the diffusion equation Eq. 4, the numerical Euler step reads

$$c_i(t_{n+1}) = c_i(t_n) \;\; - \Delta t\ (\beta + g)c_i(t_n) \;\; + \Delta t\ D\frac{2}{x_{i+1} - x_{i-1}}\left(\frac{c_{i+1}(t_n) - c_i(t_n)}{x_{i+1} - x_i} - \frac{c_i(t_n) - c_{i-1}(t_n)}{x_i - x_{i-1}}\right) + \Delta t\ \alpha\,\chi_w(x_i) \quad . \tag{S4}$$

Numerical stability requires that a Courant criterion $2D\,\Delta t \le \Delta x^2$ is satisfied.

We confirmed in selected simulations that a conventional numerical scheme using Eulerian coordinates gives identical results. Specifically, for these tests, we employed fixed spatial bins with bin centers $x_i$ and retained the advection term $v\partial_x c$ from Eq. (S1), which was discretized according to an upwind scheme, where we made use of $v_i \ge 0$. Compared to Eq. S4, the Euler step thus contains an additional advection term and reads

$$c_i(t_{n+1}) = c_i(t_n) + \text{degradation} + \text{diffusion} + \text{production} + v_i(t_n)\left(\frac{c_i(t_n) - c_{i-1}(t_n)}{x_i - x_{i-1}}\right) \quad . \tag{S5}$$

This numerical scheme with Eulerian coordinates imposes an additional criterion on the timestep $\Delta t$ for the Euler scheme $|v|\,\Delta t \le \Delta x$. If speeds $v$ become large, this criterion can be much more restrictive than the Courant criterion for the diffusion term. As a consequence, computations using Eulerian coordinates require smaller time-steps to achieve the same accuracy as computations using Lagrangian coordinates.

#### B. Image analysis

First, all 3D images were re-binned for isotropic voxel size with dimensions $2 \times 2 \times 2$ $\mu$m, then rotated and cropped in 3D with Napari [49]. Blastemas were segmented using APOC [48], trained with user-specified ground truth labels marking the tissue and the outside areas, respectively, and applied to the *Shh* channel. A constant background, corresponding to the mean intensity outside the segmented blastema, was subtracted from each channel. The blastema contained a significant number of blood cells, resulting in imaging artifacts across all channels. To address this issue, autofluorescence was acquired (499-nm illumination), and APOC segmentation was applied to this channel to identify a binary blood-vessel mask, which was subsequently excluded from all channels. Additionally, for some images, the intensity of the *Shh* channel decayed significantly along the z-axis. This artifact was corrected by normalizing the *Shh* intensity along the axis using the intensity profile calculated at the most anterior edge of the tissue, outside the possible *Shh* region.

The *Shh*, *Fgf8*, and *Dusp6* channels were denoised using a 3D Gaussian blur with a standard deviation of 10 $\mu$m (5 pixels), followed by min-max normalization. The SHH and FGF8 source regions, as well as the FGF8 signaling range region indicated by *Dusp6* mRNA in 3D, were determined as regions enclosed by the respective 50%-iso-intensity surfaces. Small disconnected components were manually removed from the volumes enclosed by these iso-intensity surfaces. The linear size of these regions were determined as the cubic roots of the volumes enclosed by the respective surfaces. Choosing different iso-intensity surfaces, i.e., at 40% or 60% relative intensity, gave the same qualitative conclusions about scaling as Fig. 4.

Blastema size was measured in 3D images along a line passing through the centers-of-mass of the *Shh* and the *Fgf8* regions, as indicated in Fig. 4A, inset. We will refer to this line as $SF$ in the following. At late stages of development and regeneration, the *Fgf8* region starts to attain a non-convex shape, and eventually splits into two separate domains. To determine the gap ($x_0$) between the tissue boundary and the boundary of the *Shh* and *Fgf8* regions in a robust manner, even for non-convex shapes of source regions, we first projected the three-dimensional binary masks along the dorsal-ventral axis onto a 2D-plane (using a logical OR-operation along the projection axis). Here, the dorsal-ventral axis was determined as the axis among all axes perpendicular to the line $SF$ that was maximally aligned to the principal axis with smallest singular value obtained by a principal-component analysis (PCA) of the binary mask of the limb bud or blastema. The outlines of these projections are shown in Fig. 4A. We then measured the distance $x_0$ between the tissue boundary and the boundary of each source region in this 2D projection along the line $SF$. For the visualization in Fig. 4, 2D-projections of different samples corresponding to the same animal size and time-point were aligned by registering the midpoint and orientation of the line segment $\overline{SF}$ connecting the centers-of-mass $S$ and $F$ of the *Shh* and *Fgf8* regions. Limb buds stages were assigned as introduced in [50].

#### C. Experimental Methods

*a. Animal breeding.* Larval axolotls (*Ambystoma mexicanum*) were maintained at the axolotl facility at CRTD TU Dresden as described [51]. Optimized axolotl (*Ambystoma mexicanum*) husbandry, breeding, metamorphosis, transgenesis and tamoxifen-mediated recombination were applied as in [51]. All procedures were performed according to the Animal Ethics Committee of the State of Saxony, Germany. Animals of the d/d genetic backgrounds with body length in the range 1.2-9 cm were used for the HCR staining. For live measurements of blastema size, axolotls of the ZRS>TFP background (tgSceI(ZRS:TFPnls-T2A-ERT2-Cre-ERT2)$^{\mathrm{Etnka}}$ [16] with body length in the range 1.2-25 cm length were used.

*b. Limb amputation and tissue collection* Axolotls were anesthetized in 0.01% benzocaine solution for approximately 20 minutes. Sleeping animals were placed on a clean Petri dish to image and measure the size of the limb. Limbs were amputated using a scalpel such that the distal quarter of the zeugopod was removed and the protruding radius and ulna were trimmed with scissors to create a flat surface. To monitor the progress of regeneration, the animals were again anesthetized in 0.01% benzocaine and imaged using a SZX16 wide field microscope with bright field illumination or UV light for the ZRS>TFP line to visualize the expression of TFP protein. Prior to tissue collection, the animals were anesthetized and limbs were imaged to record the blastema stage. For staining, limbs were cut off at the shoulder level and fixed in MEMFA (0.1 M MOPS pH 7.4, 2 mM EGTA, 1 mM $MgSO_4 \cdot 7H_2O$ and 3.7% formaldehyde) for at least 3 days at 4°C. For qPCR, only blastema and a sliver of 0.5-1 mm of tissue behind the amputation plane were collected, immediately frozen in liquid nitrogen and kept at -70°C until use.

*c. Whole-mount HCR in situ hybridization and tissue clearing.* Target transcripts were identified in the axolotl transcriptome Amex.T_v47 [52] and confirmed by BLASTX alignment to the NCBI Refseq Protein Database. HCR probe pairs were designed by the `https://probegenerator.herokuapp.com/` probe generator, which uses Bowtie2 [53] to exclude probes that hit multiple genomic regions in the 60DD axolotl genome. Probe pairs (30-40 per target transcript) were ordered as oPools from Integrated DNA Technologies. Other consumables, including hybridization, wash and amplification buffers, as well as HCR hairpins were purchased from Molecular Instruments. RNA staining was done using the whole-mount HCR protocol from [47] with the following modifications. Tissues were collected and fixed in MEMFA (3.7% formaldehyde in MEM buffer) for $>$ 3 days at 4°C. Fixative was washed out by 3 - 15 minute washes in PBST (PBS +0.1% Tween), dehydrated with 25%, 50%, and 75% MeOH in PBST all on ice and finally stored at -20°C until use (up to 2 years). After re-hydration, the permeabilization step using Proteinase K was omitted and replaced by delipidation in boric acid buffer (200 mM boric acid, 4% SDS at pH8.5) for 2-4 h at room temperature, and permeabilization in 2% TritonX-100, 0.3M glycine, 20% DMSO in PBS either at 20°C or

37°C, for 12-14h. In the detection step, the initiator pools for all target genes of interest were combined, with the concentration of each initiator pool in the hybridization solution increased to 20 nM and the reaction extended to 20 h. In the signal amplification step, Alexa-546, Alexa-647 and Alexa-750 conjugated hairpins were used at 60 nM each to detect Fgf8, Dusp6 and Shh mRNA, respectively. Both the detection and amplification steps were extended to 20-24 h. Following the final washes of the amplification step, nuclei were stained by incubation in 5xSSCT with DAPI (4,6-diamidino-2-phenylindole Sigma, #D9542) overnight at 4°C. For sample clearing, first, an equal amount of EasyIndex RI 1.52 (LifeCanvas Technologies) was added for 1 day at 4°C. Limbs were glued into a homemade vessel and covered with 100% EasyIndex for 3-10 days prior to imaging. All incubation steps were performed on a shaker or rocker. In all steps following the addition of the amplifiers, the samples were protected from light.

*d. Whole-mount imaging.* The whole-mount imaging was performed using an upright laser scanning confocal microscope Stellaris 8 (Leica) with a long range pulsed white light laser (flexible range 440-790 nm) and a pulsed 405-nm laser. For detection, LAS X software and a HC FLUOTAR L 16x/0,60 IMM CORR VISIR (11506533) Mag=16x, NA=0.6, WD=2.5mm, objective were used. The detection objective was set to match the RI of the clearing media (1.52) prior to imaging. HCR signal of Fgf8, Shh and Dusp6 was detected using the 557-nm, 752-nm and 653-nm excitation wavelength lasers. 499-nm illumination was used to detect autofluorescence. DAPI was illuminated with the 405-nm laser.

*e. qPCR.* RNA for RT-PCR was purified using RNeasy Mini Kit (Qiagen Cat. No. 74104). Reverse transcription was performed using SuperScript$^{TM}$ IV Reverse Transcriptase (Invitrogen$^{TM}$, Can. No.15387686) and Oligo dT17V as a primer. The quantitative PCR was performed using TB Green$^{(R)}$ Premix Ex Taq$^{TM}$ (Tli RNase H Plus) kit (Takara, Cat. No. RR420L).

The following primers were used:

| Target gene | Forward | Reverse |
|---|---|---|
| *Fgf8* | 829 CCGCATTAAAGGAGCCGAGA | 830 ATGAGGCTCCGTTGTTTGGT |
| *Dusp6* | 340 CCAGGACTGTCTCAAAGATTCTGC | 341 GCCTTCCGCAAGCATACATTG |
| *Shh* | 806 GCTCTGTGAAAGCAGAGAACTCG | 807 CGCTCCGTCTCTATCACGTAGAA |
| *Ptch1* | 802 CGATCGAGAAAGTGAGGGCCATA | 803 GAACTCCACTCCAATGCCAACAG |
| *Rpl4* | 185 TGAAGAACTTGAGGGTCATGG | 186 CTTGGCGTCTGCAGATTTTTT |

Analysis was performed for two biological replicates. The parallel processing of samples should make individual normalized mRNA concentrations comparable for the two animal sizes.

#### D. Statistical Methods

*a. Regression analysis.* Unweighted linear regressions by least-square-fit for slope and intercept were performed for Figs. 1C, 4D. Statistical significance of non-zero slopes in Fig. 4D was confirmed by two-sided $t$-test at confidence level $\alpha = 0.05$. Fig. 4D, left and right columns, show linear regressions against blastema width $L$ and animal size $L_a$ (snout-to-tail), respectively.

For Fig. S3, linear regressions and power-law fits were performed, to test for isometric versus allometric scaling, respectively. The power-law exponents of allometric scaling obtained here perfectly match those reported by Furukawa et al. [8]: Furukawa et al. reported exponents for the fitted allometric scaling relation between limb width and animal size of $0.90 \pm 0.06$ for fore limb and $0.91 \pm 0.07$ for hind limb, and an exponent $0.61 \pm 0.03$ for the fitted allometric scaling relationship between blastema width (late bud stage) and limb stump width (fit±SD from bootstrap), while our fits for our data gave an exponent of $0.92 \pm 0.009$ for the fitted allometric scaling relationship between fore limb width and animal length (snout-to-tail), and $0.63 \pm 0.03$ for the allometric scaling relationship between blastema width (mid/late stage) and limb width. Thus, the fitted scaling exponents almost perfectly match, although Furukawa et al. measured blastema width at the base of the blastema, while in this study (Fig. S2) blastema width was measured along a line that passes through the center of the TFP>ZRS reporter expression, parallel to the amputation plane, which is more distal towards the blastema tip compared to Furukawa, (correspondingly blastema widths are approximately 20% smaller compared to [8]). In addition to late-bud-stage blastemas, Furukawa et al. reported allometric scaling with exponent $0.43 \pm 0.07$ for blastema width as a function of limb width at the end of the first phase of blastema growth, shortly before the hypometric limb stage, when regenerated limbs had 3-4 cartilage condensations.)

*b. Partial correlation analysis.* To test for *dynamic scaling* versus *static scaling* of SHH and FGF8 morphogen gradient parameters, a partial correlation analysis was performed on the data presented in Fig. 4B as follows.

Let the observable $X$ denote a morphogen gradient parameter. The statistical factors animal size $Z = L_a$ and blastema size $Y = L$ may both influence $X$. If the null hypothesis is that $X$ exhibits *static scaling* with animal size,

$X$ should strongly correlate with $Z$, yet will be also correlated with $Y$ because $Y$ and $Z$ are correlated. To factor out the effect of $Z$, we first performed separate linear regressions

$$X_i = \alpha_X + \beta_X Z_i + \varepsilon_{X,i} \text{ and } Y_i = \alpha_Y + \beta_Y Z_i + \varepsilon_{Y,i} \quad , \tag{S6}$$

where $i = 1, \ldots, n$ enumerates the samples, and $\varepsilon_{X,i}$ and $\varepsilon_{Y,i}$ denote the respective residuals for $X$ and $Y$. Next, we perform a linear regression between these residuals to reveal effects of $Y$ on $X$ not yet explained by $Z$

$$\varepsilon_{X,i} = \alpha + \beta\, \varepsilon_{Y,i} + \varepsilon_i \quad . \tag{S7}$$

The fraction $\eta^2$ of residual variance of $X$ explained by $Y$ after factoring out $Z$ reads

$$\eta^2 = 1 - \frac{\sum_{i=1}^n \varepsilon_i^2}{\sum_{i=1}^n \varepsilon_{X,i}^2} \quad . \tag{S8}$$

The statistical significance of the effect of $Y$ on $X$ after factoring out $Z$ can be tested as follows. Let $r_{AB}$ denote the correlation coefficient between two variables $A$ and $B$ (Bravais-Pearson correlation)

$$r_{AB} = \frac{\sum_{i=1}^n (A_i - \overline{A})(B_i - \overline{B})}{\sqrt{\left(\sum_{i=1}^n \left(A_i - \overline{A}\right)^2\right)\left(\sum_{i=1}^n \left(B_i - \overline{B}\right)^2\right)}} \quad , \tag{S9}$$

where $\overline{A}$ and $\overline{B}$ denote the sample means of $A$ and $B$, respectively. The *partial correlation coefficient* $r_{XY,Z}$ between $X$ and $Y$ after factoring out $Z$ is then given by [54]

$$r_{XY,Z} = \frac{r_{XY} - r_{XZ} r_{YZ}}{\sqrt{(1 - r_{XZ}^2)(1 - r_{YZ}^2)}} \quad . \tag{S10}$$

If the null hypothesis that $Y$ has no effect on $X$ after factoring out $Z$ is true (and errors are independent and distributed normally with equal variance), the derived test-variable $t_{XY,Z}$ obeys a $t$-distribution with $n - 3$ degrees of freedom

$$t_{XY,Z} = \sqrt{n-3} \frac{r_{XY,Z}}{\sqrt{1 - r_{XY,Z}^2}} \quad . \tag{S11}$$

Now, $p$-values of a one-sided $t$-test can be readily computed. To test for the null hypothesis that a morphogen gradient parameter $X$ exhibits *dynamic scaling*, the above analysis can be repeated with the roles of $Y$ and $Z$ swapped. Results are reported in Table S3. The data reported in Fig. 4B is thus compatible with dynamic scaling of source region size $w$ and source position $x_0$ for SHH, and static scaling of source region size $w$ for FGF8. For computations, the Python package `causal-learn` was used.

#### II. SUPPLEMENTARY INFORMATION FOR MATHEMATICAL MODELING

##### A. Steady-state solutions of diffusion equation

*a. Finite system size and extended source region.* We recall Eq. 1 from the main text

$$\partial_t c(x,t) = D \partial_x^2 c(x,t) - \beta c(x,t) + \alpha \chi(x) \quad . \tag{S12}$$

We assume reflecting boundary conditions, i.e., no flux of morphogens at the system's boundaries $x = 0$ and $x = L$

$$\partial_x c(x)|_{x=0} = \partial_x c(x)|_{x=L} = 0 \quad . \tag{S13}$$

At steady state, Eq. 1 reduces to

$$0 = \lambda^2 \partial_x^2 c(x,t) - c(x,t) + \frac{\alpha}{\beta} \chi(x) \quad , \tag{S14}$$

where $\lambda = (D/\beta)^{1/2}$ denotes the pattern length-scale as introduced in the main text. The Green's function for the differential operator $\lambda^2\partial^2 - 1$ with reflective boundary conditions Eq. S13 reads

$$G(x,s) = \frac{1}{\lambda \sinh \frac{L}{\lambda}} \left[ \cosh\left(\frac{L-s}{\lambda}\right) \cosh\left(\frac{x}{\lambda}\right) \Theta(s-x) + \cosh\left(\frac{s}{\lambda}\right) \cosh\left(\frac{L-x}{\lambda}\right) \Theta(x-s) \right] \quad , \tag{S15}$$

where $\Theta(x)$ denotes the Heaviside step function. Thus, the steady state solution $c^*$ of Eq. (1) can be written for an arbitrary source region described by a shape function $\chi(x)$ as

$$c^*(x) = \frac{\alpha}{\beta} \int_0^L G(x,s)\ \chi(s)\ ds \quad . \tag{S16}$$

For the source function introduced in the main text

$$\chi_w(x) = \Theta(w + x_0 - x)\Theta(x - x_0) \quad , \tag{S17}$$

which takes the constant value 1 inside the interval $[x_0, x_0 + w]$ and zero outside, the steady state solution $c^*$ reads

$$c^*(x) = \frac{\alpha}{\beta \sinh \frac{L}{\lambda}} \begin{cases} 2 \sinh \frac{w}{2\lambda} \cosh \frac{2L-w-2x_0}{2\lambda} \cosh \frac{x}{\lambda}, & x \in [0, x_0) \\ \sinh \frac{L}{\lambda} - \sinh \frac{L-w-x_0}{\lambda} \cosh \frac{x}{\lambda} - \sinh \frac{x_0}{\lambda} \cosh \frac{L-x}{\lambda}, & x \in [x_0, x_0 + w] \\ 2 \sinh \frac{w}{2\lambda} \cosh \frac{w+2x_0}{2\lambda} \cosh \frac{L-x}{\lambda}. & x \in (x_0 + w, L] \end{cases} \tag{S18}$$

*b. Point source region.* To facilitate analytical analysis of growth arrest and morphogen gradient scaling as a function of key model parameters (diffusion coefficient $D$, degradation rate $\beta$, and total production rate $j = \alpha w$), we state the steady-state solution of Eq. (S18) for a point morphogen source region located at $x = x_0$, corresponding to the limit $w \to 0$ with $j = \alpha w = \text{const}$.

$$c^*(x) = \frac{\alpha w}{\sqrt{D\beta} \sinh \frac{L}{\lambda}} \begin{cases} \cosh \frac{L-x_0}{\lambda} \cosh \frac{x}{\lambda}, & x \in [0, x_0] \\ \cosh \frac{x_0}{\lambda} \cosh \frac{L-x}{\lambda}. & x \in (x_0, L] \end{cases} \tag{S19}$$

*c. Point source region at system boundary.* For sake of completeness, we additionally state the steady-state solution of Eq. 1 for a point source located the the system boundary $x_0 = 0$

$$c^*(x) = \frac{\alpha w}{\sqrt{D\beta} \sinh \frac{L}{\lambda}} \cosh \frac{L-x}{\lambda} \quad . \tag{S20}$$

As a technical point, the no-flux boundary condition S13 needs to be modified for a point source located at the system boundary such that the diffusive flux equals morphogen production, $-D\partial_x c_{|x=0} = j$.

*d. Point source region in an unbounded system.* In the limit of an unbounded system with $L \to +\infty$, the steady-state solution Eq. S20 further simplifies to

$$c^*(x) = \frac{\alpha w}{\sqrt{D\beta}} \exp\left(-\frac{x}{\lambda}\right) \quad , \tag{S21}$$

corresponding to Eq. (3) in the main text.

*e. Point source region with absorbing boundary.* Finally, one could assume absorbing boundary conditions, i.e., the system's boundaries $x = 0$ and $x = L$ act as perfect sinks, and morphogen concentrations must vanish there

$$c(0) = c(L) = 0 \quad . \tag{S22}$$

For this case, the steady-state solution for Eq. 1 for a point source located at $x_0$ reads

$$c^*(x) = \begin{cases} \dfrac{\alpha w \lambda}{D} \dfrac{\sinh\left(\frac{L-x_0}{\lambda}\right) \sinh\left(\frac{x}{\lambda}\right)}{\sinh\left(\frac{L}{\lambda}\right)}, & x \in (0, x_0] \\ \dfrac{\alpha w \lambda}{D} \dfrac{\sinh\left(\frac{x_0}{\lambda}\right) \sinh\left(\frac{L-x}{\lambda}\right)}{\sinh\left(\frac{L}{\lambda}\right)}, & x \in (x_0, L) \end{cases} \quad . \tag{S23}$$

##### B. Oppositely-oriented morphogen gradients at left and right system boundaries

As mentioned in main text, the degradation-diffusion model for morphogen dynamics Eq. (1) can be generalized in a straight-forward manner to the case of two opposing morphogen gradients with respective concentration fields $s(x,t)$ and $f(x,t)$, which can be explicitly written as

$$\partial_t s(x,t) = D\,\partial_x^2 s(x,t) - \beta\, s(x,t) + \alpha\,\chi_s(x) \quad , \tag{S24}$$

$$\partial_t f(x,t) = D\,\partial_x^2 f(x,t) - \beta\, f(x,t) + \alpha\,\chi_f(x) \quad , \tag{S25}$$

where, for simplicity, the source regions are also mirror-symmetric, $\chi_s(x) = \chi_f(L-x)$.

##### C. Growth rules define the final system size as a function of model parameters

*a. One-morphogen growth rule.* In the case of the one-morphogen growth rule, growth stops when the growth threshold $\Theta$ equals the morphogen gradient amplitude $A$. The final system size $L^*$ is thus determined by the condition

$$L^*_{\text{one}} : A := \max_{[0,L^*]} c(x) \stackrel{!}{=} \Theta \quad . \tag{S26}$$

In the limit case of a point morphogen source Eq. S20, the final system size can be written as

$$L^*_{\text{one}} = \lambda \ \text{arcoth}\left(\frac{\Theta\sqrt{D\beta}}{\alpha w}\right) \quad . \tag{S27}$$

The function $\text{arcoth}\left(\frac{\Theta\sqrt{D\beta}}{\alpha w}\right)$ is defined only for $\frac{\Theta\sqrt{D\beta}}{\alpha w} > 1$, which gives the critical growth threshold $\Theta_c$ for growth arrest for the one-morphogen growth rule as

$$\Theta_c = \frac{\alpha w}{\sqrt{D\beta}} \quad . \tag{S28}$$

Growth arrest occurs for $\Theta > \Theta_c$.

*b. Two-morphogen growth rule.* For the two-morphogen growth rule, growth stops when the growth threshold $\Theta$ equals the morphogen concentration at the intersection of the morphogen. For simplicity, we consider the symmetrical case, where both morphogens have identical parameters, but their source regions are located at opposite boundaries of the system. In this case, morphogen gradients intersect at the midpoint of the system; therefore, the final system size $L^*$ is determined by

$$L^*_{\text{two}} : c\left(\frac{L^*}{2}\right) \stackrel{!}{=} \Theta \quad . \tag{S29}$$

In the limit case of a point morphogen source, Eq. S20, we have

$$c\left(\frac{L}{2}\right) = \frac{aw}{\sqrt{D\beta}}\frac{\cosh(L/2\lambda)}{\sinh(L/\lambda)} = \frac{\Theta_c}{2}\,\text{csch}\left(\frac{L}{2\lambda}\right) \quad . \tag{S30}$$

Thus, in case of growth arrest, we have

$$\frac{\Theta_c}{2\Theta} \stackrel{!}{=} \sinh\left(\frac{L}{2\lambda}\right) \quad , \tag{S31}$$

and the final system size reads

$$L^*_{\text{two}} = 2\lambda \ \text{arsinh}\left(\frac{\alpha w}{2\Theta\sqrt{D\beta}}\right) = 2\lambda \ \text{arsinh}\left(\frac{\Theta_c}{2\Theta}\right) \quad . \tag{S32}$$

From Eq. S30, we conclude

- For static scaling (Fig. 3D), growth arrest occurs for any $\Theta$ (as $\Theta_c$ and $\lambda$ stay constant during growth, and thus $c(L/2)$ monotonically decreases to zero for $L \to \infty$).
- For dynamic scaling without scaling of $\lambda$ (Fig. 3C, first column), growth arrest also occurs for any $\Theta$ (as $\Theta_c \sim L$, $\lambda$ stays constant, and $c(L/2) \sim \xi\,\text{csch}\,\xi$ monotonically decreases to zero as function of $\xi = L/2\lambda \to \infty$).
- For dynamic $D$-scaling (Fig. 3C, second column), growth does not arrest (as $\Theta_c$ and $L/2\lambda$ stay constant during growth, hence by Eq. S30 also $c(L/2)$) stays constant. More generally, Eq. S18 implies that $c(L/2)$ stays constant in this scenario also for a source of finite width with $w \sim L$ and $x_0 \sim L$.

*c. Mixed scaling variants for D-scaling.* Fig. S11C-E numerically analyzed variants of mixed scaling, where different morphogen parameters for the two morphogen profiles exhibit either static or dynamic scaling. We give an analytical argument why proportional growth is expected even for these mixed scaling variants in the limit case of a point morphogen source at the system's boundary if $D$-scaling is assumed. We introduce the dimensionless quantity $\Lambda = L/L_a$. The dependence of $\Theta_c$ and $L/\lambda$ on $L$ can now be expressed as a dependence on $\Lambda$, see the table below.

| **Parameter** | **Variants** | | | |
|---|---|---|---|---|
| $w$ | static | dynamic | static | dynamic |
| $\lambda$ | $D$-scaling: static | | $D$-scaling: dynamic | |
| $\Theta_c$ | $\sim 1$ | $\sim \Lambda$ | $\sim \Lambda^{-1}$ | $\sim 1$ |
| $L/\lambda$ | $\sim \Lambda$ | | $\sim 1$ | |

According to Eq. S20, also the dependence of the morphogen profile $c^*(x)$ on the growing system size $L$ can be expressed as a dependence on $\Lambda$, for which we write $c_\Lambda(x/L)$. In the case of two oppositely-oriented morphogen profiles $s(x)$ and $f(x)$, possibly with different morphogen parameters and different mixed scaling dependencies, we can express their dependence on $L$ through $\Lambda$ as $s_\Lambda(x/L)$ and $f_\Lambda(x/L)$, respectively. If growth arrest occurs at a final system size $L^*$, the growth zone will have shrunken to a single point $x^*$. Hence, according to the two-morphogen growth rule, we must have

$$s_{\Lambda^*}(x^*/L^*) \stackrel{!}{=} \Theta \text{ and } f_{\Lambda^*}(x^*/L^*) \stackrel{!}{=} \Theta \quad , \tag{S33}$$

where $\Lambda^* = L^*/L_a$. Solving for $x^*/L^*$, we obtain the condition

$$s_{\Lambda^*}^{-1}(\Theta) = f_{\Lambda^*}^{-1}(\Theta) \quad . \tag{S34}$$

If growth arrest occurs, the transcendental equation Eq. S34 has a solution $\Lambda^*$. But this implies $L^* = \Lambda^* L_a$, i.e., if there is growth arrest, there is also proportional growth. The above argument still holds if for each morphogen different growth thresholds $\Theta_s$ and $\Theta_f$ are assumed.

For sake of explicitness, we state Eq. S34 for an example scenario where

- the morphogen $s(x)$ exhibits *dynamic* source scaling ($w_s$) and *static* $D$-scaling of its pattern length scale $\lambda_s$, which

$$s_\Lambda\left(\frac{x}{L}\right) = \Theta_{c,s}\frac{\cosh\left[\frac{L}{\lambda_s}\left(1-\frac{x}{L}\right)\right]}{\sinh\left(\frac{L}{\lambda_s}\right)} = b_s\Lambda\frac{\cosh\left[a_s\Lambda\left(1-\frac{x}{L}\right)\right]}{\sinh\left(a_s\Lambda\right)} \quad , \tag{S35}$$

  where we used $L/\lambda_s = a_s\Lambda$, $\Theta_{c,s} = b_s\Lambda$ with constants of proportionality $a_s$, $b_s$ (see table above), and

- the morphogen $f(x)$ exhibits *static* source scaling ($w_f$) and *dynamic* $D$-scaling of its pattern length scale $\lambda_f$, which gives

$$f_\Lambda\left(\frac{x}{L}\right) = \Theta_{c,f}\frac{\cosh\left(\frac{L}{\lambda_f}\frac{x}{L}\right)}{\sinh\left(\frac{L}{\lambda_f}\right)} = b_f\Lambda^{-1}\frac{\cosh\left(a_f\frac{x}{L}\right)}{\sinh\left(a_f\right)} \quad , \tag{S36}$$

  where we used $L/\lambda_f = a_f$, $\Theta_{c,f} = b_f\Lambda^{-1}$ with constants $a_f$, $b_f$.

At growth arrest, the condition for $\Lambda^*$ reads

$$1 - \frac{1}{a_s\Lambda^*}\operatorname{arccosh}\left[\frac{\Theta_s}{b_s\Lambda^*}\sinh\left(a_s\Lambda^*\right)\right] \stackrel{!}{=} a_f^{-1}\operatorname{arccosh}\left[\frac{\Theta_f\Lambda^*}{b_f}\sinh a_f\right] \quad . \tag{S37}$$

For suitable parameters, this equation has a unique solution $\Lambda^*$ independent of other length-scales, hence $L^* \sim L_a$. It can be shown that the same argument still holds true in the general case of a source of finite width $w > 0$ and general position $x_0$, provided both source parameters scale (either dynamically or statically). This is consistent with the numerical findings shown in Fig. S7C-E.

*d. Re-scaling of units.* In the limit case of a point source with $w \ll \lambda$, it is possible to non-dimensionalize our model as follows

$$t \to \tilde{t} = \beta\, t\,, \tag{S38}$$

$$x \to \tilde{x} = x/\lambda\,, \tag{S39}$$

$$c \to \tilde{c} = c/\Theta_c\,. \tag{S40}$$

Eqs. (S19, S20, S21, S23, S27, S28, S32) above can be rewritten in terms of these re-scaled quantities. For example, Eq. S19 can be rewritten as

$$\tilde{c}^* = \mathrm{csch}\tilde{L} \begin{cases} \cosh\left(\tilde{L} - \tilde{x}_0\right)\cosh\tilde{x}, & \tilde{x} \in [0, \tilde{x}_0] \\ \cosh\left(\tilde{L} - \tilde{x}\right)\cosh\tilde{x}_0, & \tilde{x} \in (\tilde{x}_0, \tilde{L}] \end{cases} . \tag{S41}$$

Note that the decay length $\lambda$ will change dynamically for the dynamic scaling scenario, which impacts system behavior.

#### D. Mutual feedback between morphogens

In the main text, we introduced feedback between the two morphogens, where a morphogen can only be produced inside a competent zone at the boundary with width $w_{\text{competent}}$ if the other morphogen exceeds a fixed feedback threshold, $\Theta_{\text{fb}}$. To obtain the phase diagram Fig. 5B, we define a separatrix that separates a regime dominated by the growth threshold $\Theta$ and a regime dominated by the feedback threshold $\Theta_{\text{fb}}$. Specifically, the separatrix is defined by the condition that both thresholds can be reached simultaneously. Accordingly, the final source position $x_0^*$ is both strictly positive and strictly smaller than $w_{\text{competent}}$.

In our minimal model, growth arrests only if the growth threshold $\Theta$ has been reached. For the two-morphogen growth rule, this occurs at the midpoint of system, see also Eq. S29. In this case, the source width equals $w_{\text{competent}} - x_0^*$. Hence, assuming that the growth threshold is reached outside the source region and using Eqs. S18 and S29, the growth threshold can be explicitly written as

$$\Theta = \frac{2\alpha}{\beta \sinh\left(\frac{L^*}{\lambda}\right)} \sinh\left(\frac{w_{\text{comp.}} - x_0^*}{2\lambda}\right) \cosh\left(\frac{2w_{\text{comp.}} + x_0^*}{2\lambda}\right) \cosh\left(\frac{L^*}{2\lambda}\right) \quad . \tag{S42}$$

For the case of symmetrical morphogens at two opposing sides of the system, we can also state the feedback threshold at the moment of growth arrest as the concentration at the position $x = L^* - x_0^*$, which can be written as

$$\Theta_{\text{fb}} = \frac{2\alpha}{\beta \sinh\left(\frac{L^*}{\lambda}\right)} \sinh\left(\frac{w_{\text{comp.}} - x_0^*}{2\lambda}\right) \cosh\left(\frac{2w_{\text{comp.}} + x_0^*}{2\lambda}\right) \cosh\left(\frac{x_0^*}{\lambda}\right) \quad . \tag{S43}$$

Assuming the critical situation, where $x_0^* = 0$ and varying the final system size $L^*$ according to Eq. S42 and Eq. S43, we obtain the separatrix between the two regimes as shown in Fig. 5B.
